# Structural relationships between the SLC12 cotransporter C-terminal domain, universal stress proteins and adenylating enzymes

**DOI:** 10.64898/2026.11.09.765458

**Authors:** Edina Rosta, Katharina L. Dürr

## Abstract

Several SLC12 cotransporters bind ATP in a cytoplasmic C-terminal domain (CTD). We analysed 16,192 CTD-containing SLC12 sequences and compared 42,845 representative structures to examine the relationship between this nucleotide site, universal stress proteins (USPs) and adenylating enzymes. In phosphorylated NKCC1, D891 and D927 lie near the ATP phosphate chain, while R801 and K889 form a basic phosphate-binding clamp. NKCC1 retains this arrangement, whereas KCC paralogs replace or lose several corresponding groups. In structural comparisons with USP and adenylating-enzyme references, both CTD lobes score between universal stress proteins and EC-annotated domains, and the two lobes form separate lineages that include bacterial transporters. Lobe-specific and nucleotide-frame comparisons place ATP in the corresponding structural region used for α-phosphate chemistry by catalytic members of this fold. Published work on UspG provides a precedent for protein-bound AMP in a natural USP. In the examined SLC12 structures and USP models, candidate protein hydroxyls lack an in-line geometry for attack at ATP Pα. Protein adenylation would require a change in protein or nucleotide geometry, or a different acceptor. These results define a structural basis for testing conditional protein adenylation at SLC12 nucleotide sites. The required evidence is a covalent protein–AMP adduct, identification of its acceptor and measurement of the expected pyrophosphate coproduct, with soluble nucleotide analysis to distinguish accompanying ATP reactions.

**Significance:** SLC12 cotransporters control cellular chloride balance. Several family members bind adenine nucleotides in their cytoplasmic domains. Their ATP-binding lobe resembles universal stress proteins (USPs) and shares a broader fold with adenylating enzymes. We quantify these structural relationships and identify the changes in geometry required for candidate protein hydroxyls to attack ATP. Reported protein-bound AMP in UspG supplies a biochemical precedent in a natural USP. Together, these observations motivate a direct test for covalent AMP attachment at SLC12 nucleotide sites, with protein–AMP and pyrophosphate as the expected products.

## Introduction

The cation-chloride cotransporter family (SLC12) controls intracellular Cl^−^ homeostasis and contributes to inhibitory neurotransmission, renal salt handling and epithelial fluid balance (1, 2). NKCC1 (SLC12A2), NKCC2 (SLC12A1), NCC (SLC12A3) and KCC1–4 (SLC12A4–7) contain a cytoplasmic C-terminal regulatory domain (CTD). Phosphorylation through the WNK/SPAK/OSR1 pathway regulates their transport activity, activating NKCCs and inhibiting KCCs (3, 4). Several family members also bind adenine nucleotides in the CTD, providing a potential connection between the nucleotide state and transporter regulation (5, 6, 7, 8).

Structural studies have defined the CTD architecture in archaeal CCCs, a eukaryotic KCC regulatory domain and full-length KCC and NKCC proteins (9, 10, 11, 12, 13). Warmuth et al. identified a distant structural relationship between the archaeal CTD and universal stress proteins (USPs) (9). Chi et al. subsequently identified TeaD as the closest DALI match to the KCC1 nucleotide-binding lobe and proposed a possible role for conserved acidic residues in ATP hydrolysis (5). The KCC, NCC and phosphorylated NKCC1 studies also support structural or allosteric roles for bound nucleotides (5, 6, 7, 8). In phosphorylated NKCC1, ATP–Mg^2+^ stabilises otherwise dynamic CTD elements (8). These observations establish the structural relationship and motivate a quantitative comparison of the nucleotide site.

The evolutionary HUP framework links USP, HIGH-containing class-I aminoacyl-tRNA synthetase and PP-ATPase domain lineages (14). SCOPe places selected members of these lineages within the adenine nucleotide α-hydrolase-like fold, c.26 (15). Within this fold, USPs and the catalytic domain of β-lactam synthetase belong to the c.26.2 superfamily, whereas class-I aminoacyl-tRNA synthetase catalytic domains belong to the distinct c.26.1 superfamily (Figure 1). β-Lactam synthetase and class-I aminoacyl-tRNA synthetases cleave the ATP α– β bond during substrate adenylation, releasing pyrophosphate (PPi) (16, 17). HUP describes the broader evolutionary relationship; the SCOPe hierarchy distinguishes the fold, superfamilies and families used here.

**Figure 1.**
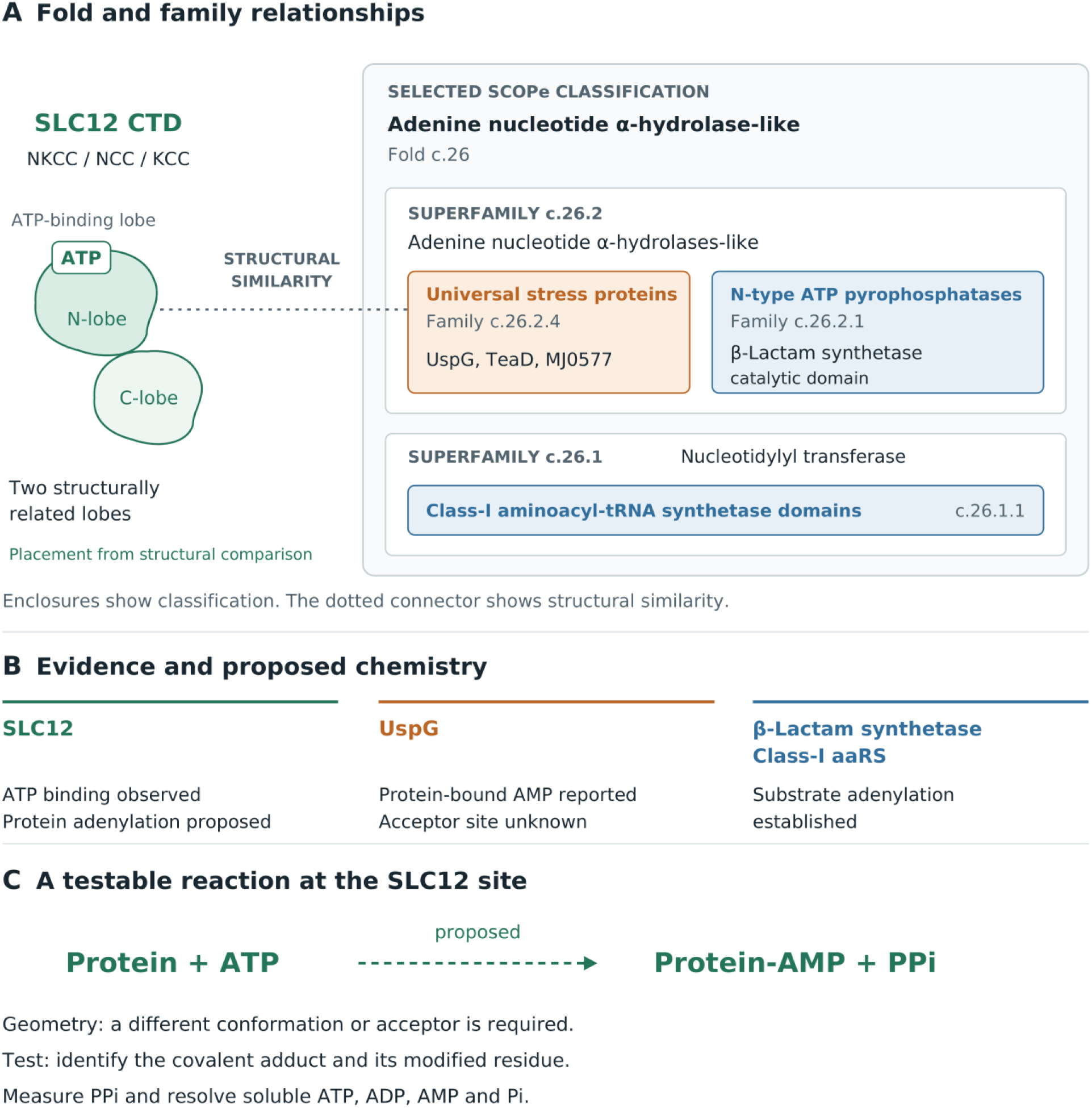
Structural relationships and the SLC12 protein-adenylation hypothesis. (A) Selected SCOPe families within the adenine nucleotide α-hydrolase-like fold, c.26 (15). USPs and N-type ATP pyrophosphatases, including the catalytic domain of β-lactam synthetase, belong to superfamily c.26.2; class-I aminoacyl-tRNA synthetase catalytic domains belong to c.26.1. The dotted SLC12–USP connector denotes structural similarity. The SLC12 CTD contains two structurally related lobes, with the illustrated ATP site in the N-lobe. The enclosures show classification and the shapes show schematic domain architecture. (B) Evidence distinguishes observed ATP binding in SLC12, reported covalent protein-bound AMP in UspG and established substrate adenylation in the enzyme references (5, 16, 17, 21). The UspG acceptor remains unassigned. (C) The proposed SLC12 reaction predicts protein–AMP and PPi. Testing it requires identification of the adduct and its acceptor, together with inorganic-phosphate and soluble-nucleotide analysis. The candidate hydroxyls examined here require a change in protein or nucleotide geometry, or another acceptor.

Sequence-domain annotations define the comparison sets used in this study. The SLC12 CTD has a dedicated InterPro annotation, IPR018491; the broader IPR014729 annotation supplied the principal structural survey, with SLC12 proteins retrieved separately (18). Earlier analyses of functional divergence within HUP and ATP-PPase domains provide context for interpreting this diverse comparison set (19, 20). We use structural similarity to place SLC12 relative to the sampled USP and enzyme references. The opening schematic distinguishes this structural comparison from the selected SCOPe classifications (Figure 1).

We quantify the structural relationship between SLC12 CTDs, USPs and adenylating enzymes, then examine the geometric requirements for protein adenylation at the transporter nucleotide site. Sequence comparison of 16,192 CTD-containing proteins defines the retention of the D891/D927/R801/K889 pocket arrangement in phosphorylated NKCC1. A survey of 42,845 structural representatives locates SLC12 relative to USP and enzyme references. Lobe-specific and nucleotide-frame comparisons then identify the position of ATP and the geometric changes required for candidate protein acceptors.

## Results

### ATP pocket architecture in phosphorylated NKCC1

The phosphorylated NKCC1–ATP–Mg^2+^ structure, 9C0E (K289N/A492E construct with furosemide, 2.7 Å), provides an experimental view of the nucleotide pocket (8). D891 OD1 lies 3.9 Å from ATP Pβ and D927 OD1 lies 5.2 Å from Pγ (OD2 4.8 Å) in chain A of the analysed coordinates (Figure 2). R801 and K889 contact the phosphate chain and form a basic clamp, as described by Zhao et al. (8). Mg^2+^ contacts the ATP phosphates; the two Asp residues lie outside its first coordination shell. These contacts define the acidic and basic groups surrounding the bound nucleotide.

**Figure 2.**
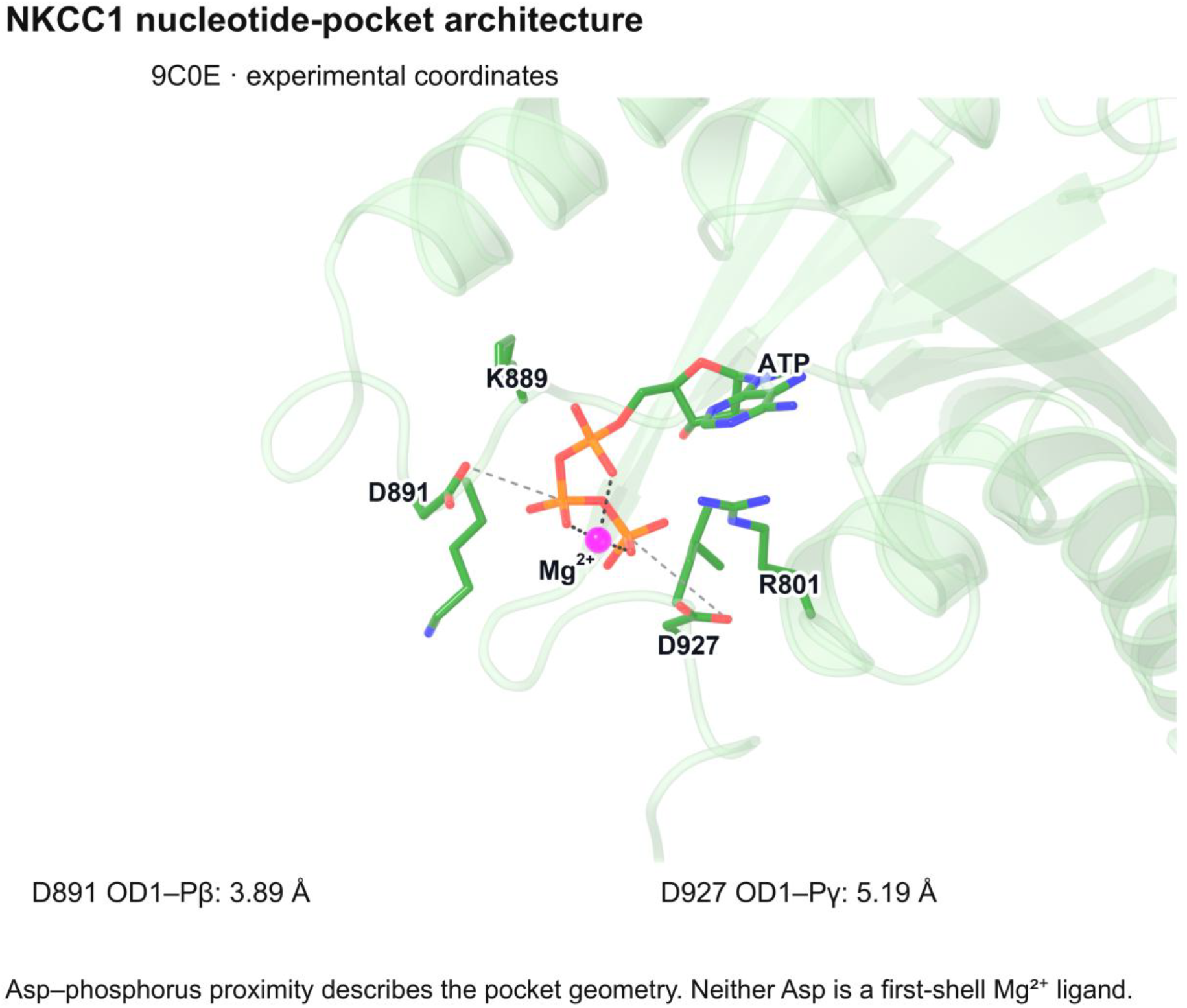
Experimental NKCC1 nucleotide-pocket architecture in 9C0E (8). D891 OD1 lies 3.89 Å from ATP Pβ and D927 OD1 lies 5.19 Å from Pγ (OD2 4.78 Å) in chain A of the saved coordinates. R801 and K889 form a basic phosphate-binding clamp. The modelled Mg^2+^ contacts nucleotide oxygens, while the two Asp residues lie outside its first coordination shell. The distances describe the observed arrangement.

Across the family alignment, strict Asp retention among non-gap entries is 49.7% at the KCC1 D804-corresponding position, 19.0% at NKCC1 D891 and 53.1% at D927. D891 is the most variable of these positions. We distinguish acidic D/E pairs from pairs containing N/Q and from other residue combinations (Table 1). These classes describe the charge and functional groups available at the two positions; metal coordination depends on their three-dimensional arrangement.

**Table 1.** Chemical classes at the positions corresponding to NKCC1 D891 and D927. The definitions apply to pairs of known amino acids. Gaps and ambiguous residues are unassigned. The classes describe functional groups at the two positions; they do not specify a metal-coordination geometry.

| Class | Residue pair | Chemical groups |
|---|---|---|
| I | D/E at both positions | Two carboxylate side chains |
| II | D/E/N/Q at both positions, with N/Q at one or both | At least one amide side chain; two potential oxygen-donor side chains |
| III | A known amino acid outside D/E/N/Q at either position | Other combinations of side-chain functional groups |

Asp retention differs among the sampled lineages (Figure S1). Both curated mammalian NKCC1 orthologues, human and mouse, retain both Asp residues. In the broader keyword-assigned groups, insects retain both in 37.7% of sequences, plants retain D927 alone in 90.8%, and fungi retain neither in 98.2%. The 10,898 sequences assigned to “Other” comprise 67.3% of the full alignment. The distributions therefore describe the sampled groups and their paralog composition; they do not measure evolutionary selection.

### KCC paralogs have remodelled the ATP binding pocket

Sequence and AlphaFold 3 comparisons identify coordinated changes in KCCs. KCC2 carries Asn at the D891-corresponding position and has no residue at the D927-corresponding position, which is a gap in all reviewed KCC sequences. It carries Pro at the K889-corresponding position. The R801 position is not conserved in KCCs; depending on the alignment, it holds His, Tyr or a gap. Pro removes the Lys side-chain amine and changes the local backbone geometry. Across reviewed paralogs, the W892-corresponding aromatic residue is broadly conserved, while R801 and K889 are retained in NKCC/NCC but not in KCCs. All reviewed KCCs carry Lys at the R921-corresponding position. These substitutions and deletions remodel several parts of the phosphate-binding surface.

In the five KCC2 AlphaFold 3 models used for this comparison, the nearest Ser/Thr hydroxyl, Thr787 OG1, lies 6.9–8.3 Å from ATP Pγ across both protomers. This comparison evaluates candidate protein acceptors; water-mediated hydrolysis requires a separate assessment of water occupancy and reaction geometry. We use the sequence alignment to establish residue identities and the predictions to illustrate their possible spatial arrangement. Experimental NKCC1 9C0E and KCC1 7AIP have different metal sites (Figure S2). They differ in paralog and in modelled phosphorylation (9C0E pThr212, pThr217 and pThr230; 7AIP pSer734 and pSer916), so the comparison cannot isolate a phosphorylation-driven motion.

### SLC12 CTD lobes lie between USPs and the enzyme references

We compared fold domains from 42,845 structural representatives with five references: the USPs MJ0577 (1MJH) and TeaD (3HGM), the archaeal CCC CTD (3G40), and the catalytic domains of β-lactam synthetase (1MB9) and glutamyl-tRNA synthetase (1J09) (9, 16, 17, 22, 23). The SLC12 CTD and the archaeal CCC CTD each contain two lobes of the fold. In whole-protein alignments, the five references did not all align to the same SLC12 lobe in 377 of 627 entries. We used each lobe as a separate query and represented the archaeal CCC by its two lobes (Methods).

Median ratios of mean Z to the USP/CCC references over mean Z to the enzyme references were 2.46 for USP domains, 1.76 for SLC12 N-lobes, 1.66 for SLC12 C-lobes and 1.22 for EC-annotated domains (Figure 3A). SLC12 N-lobes scored above EC-annotated domains with a common-language effect size of 0.948 (95% CI 0.943–0.951), and C-lobes with 0.923 (0.916–0.930). Both lobes scored below USP domains (0.081 and 0.057). The separation from the EC-annotated group persisted within domain-length quintiles and across seven scoring rules (Figure 3B, C). Without the archaeal CCC references, the effect sizes were 0.920 for N-lobes and 0.813 for C-lobes. The score is a ratio, and the difference between SLC12 and USP domains comes from its numerator. Median Z to the two enzyme references was 6.0 for USP domains, 5.7 for SLC12 N-lobes and 5.1 for C-lobes, so neither SLC12 lobe is closer to those references than USP domains are. Compared with the EC-annotated domains themselves rather than with the references, USP domains are the closer group (median best LDDT 0.65, against 0.58 for N-lobes and 0.55 for C-lobes). This ranking holds for the EC-annotated group as labelled; restricted to the enzyme-like core of that group, the three groups are equidistant (median best LDDT 0.52). A lower score on this axis records weaker similarity to the USP and CCC references, not greater similarity to enzymes.

**Figure 3.**
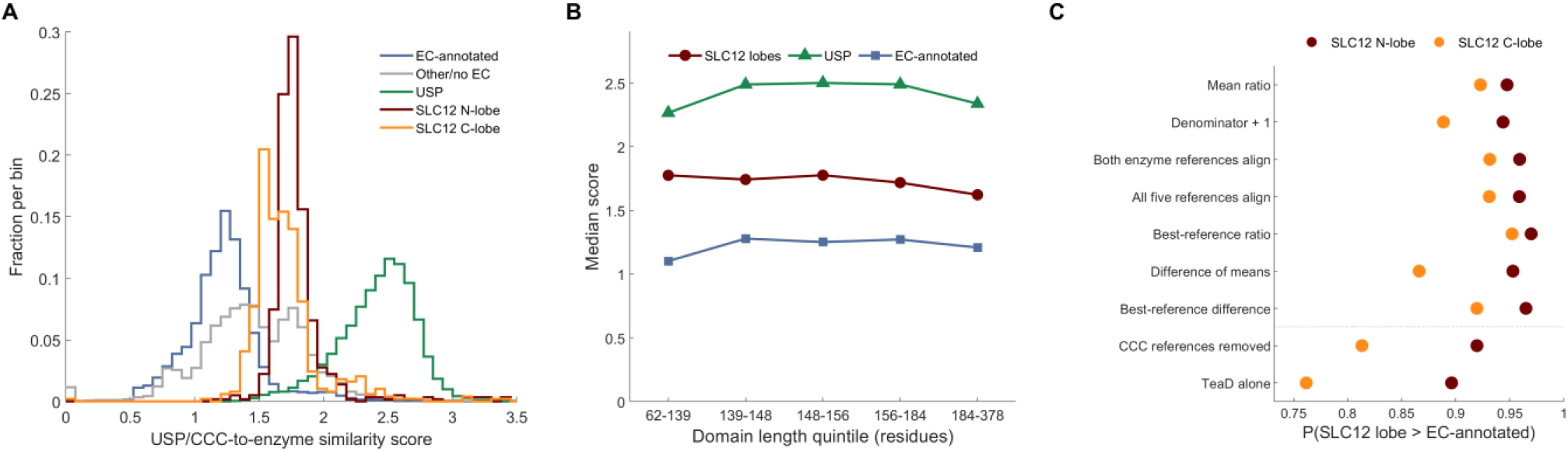
DALI similarity of fold domains to USP/CCC and enzyme references. Each SLC12 CTD lobe and each lobe of tandem USPs was a separate query. The archaeal CCC reference was split into its two lobes and contributes its higher-scoring lobe; the enzyme references are their SCOPe catalytic domains. (A) Distributions of mean Z to MJ0577, TeaD and the archaeal CCC lobe divided by mean Z to the β-lactam synthetase and glutamyl-tRNA synthetase domains, for 46,367 domains (45,378 with a defined score). Histograms use 0.075-wide bins; 220 scores above 3.5 lie outside the displayed range. (B) Median scores within quintiles of domain length defined from SLC12 lobes. (C) Common-language effect sizes for SLC12 N- and C-lobes against EC-annotated domains, for seven scoring rules and two reference-removal controls, with ties counted at half weight. A lower score records weaker similarity to the USP and CCC references; it does not place a domain closer to the enzyme references.

Each SLC12 lobe matched the corresponding archaeal CCC lobe: 534 of 585 N-lobes scored higher against the 3G40 N-lobe and 4 scored equally, and 495 of 516 C-lobes against the 3G40 C-lobe. A three-cluster partition placed 91% of USP domains, 44% of SLC12 N-lobes, 4% of SLC12 C-lobes and 2% of EC-annotated domains in one cluster. The value of k was selected for agreement with the annotation groups, so this partition is a label-informed summary.

A structure-based tree of 1,977 representative domains places the SLC12 lobes within the fold (Figure 4). Of 202 SLC12 N-lobes, 199 form a clade that contains no other domain, and 195 of 196 C-lobes form a separate clade of 196 tips; the two clades are not sisters. Each clade holds the bacterial SLC12 lobes (14 of 14 N-lobes and 13 of 13 C-lobes) and the experimental NKCC1 and KCC1 lobes, and the archaeal CCC C-lobe falls with the SLC12 C-lobes. Both lobe clades sit inside a group of 1,006 tips that holds 509 of 542 USP domains and 41 of 835 domains from the enzyme, ETF and photolyase families. Trees from two other structural distances recover the N-lobe clade, the archaeal C-lobe grouping and the bacterial lobes; one splits the C-lobes into two clades, and neither recovers the 1,006-tip group (Table S4).

**Figure 4.**
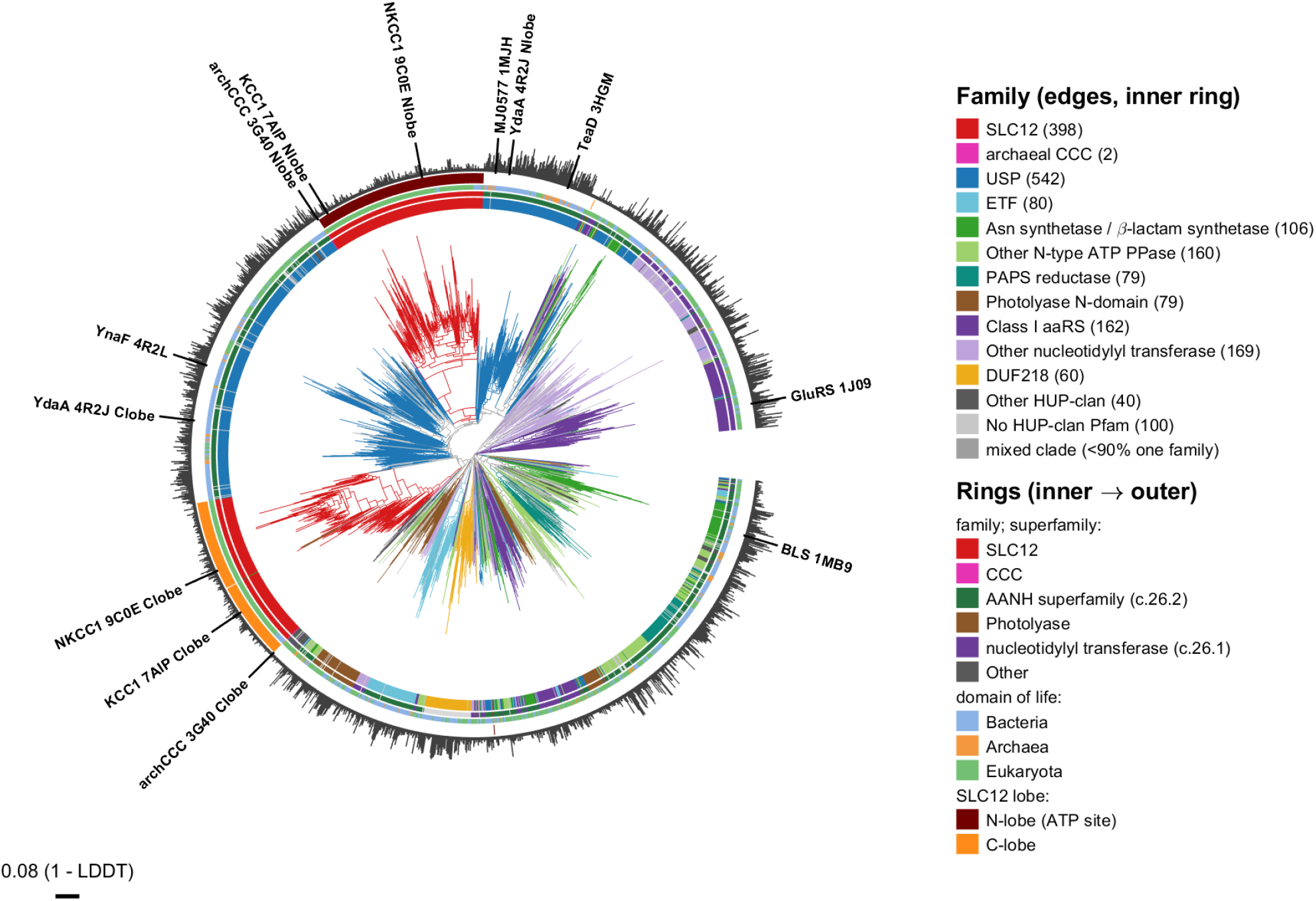
Structure-based relationships among the SLC12 CTD lobes and the comparison set. The tree contains 1,977 representative fold domains from structural clusters of the survey, including both lobes of 194 SLC12 proteins and 129 tandem USPs, and was built by balanced minimum evolution from Foldseek 1 − LDDT distances (Methods). The set comes from the InterPro-based survey and is not confined to one SCOPe fold: the USP, class-I aminoacyl-tRNA synthetase, asparagine synthetase and β-lactam synthetase, electron transfer flavoprotein and PAPS reductase families sit in the adenine nucleotide α-hydrolase-like fold c.26, while the photolyase N-terminal domains sit in the separate fold c.28 (15). Edge colour and the inner ring show the family. The next rings show superfamily, domain of life and SLC12 lobe. Outer bars give the number of surveyed domains placed on each representative. Experimental domains are labelled. SLC12 N-lobes and C-lobes form separate clades, and the archaeal CCC C-lobe is the sister of the SLC12 C-lobe clade. Across the sampled queries the median best-match LDDT of an N-lobe was higher to a USP domain than to a C-lobe, 0.587 against 0.557, and 0.575 when the USP target pool is cut to the size of the C-lobe pool. The matching C-lobe comparison falls within the sampling margin and is not reported. The root is placed on the branch that best separates the nucleotidylyl-transferase superfamily and is used for display.

### The SLC12 nucleotide site corresponds to an adenylating site in the fold

β-Lactam synthetase supplies a direct structural reference for α-phosphate chemistry. Its substrate acyl-adenylate intermediate is captured with PPi still bound in 1MBZ (16). The covalently attached oxygen belongs to the small-molecule substrate. Class-I aminoacyl-tRNA synthetases provide a second example of ATP α–β cleavage during amino-acid activation (17). Figure 5 compares these experimental nucleotide sites with NKCC1.

**Figure 5.**
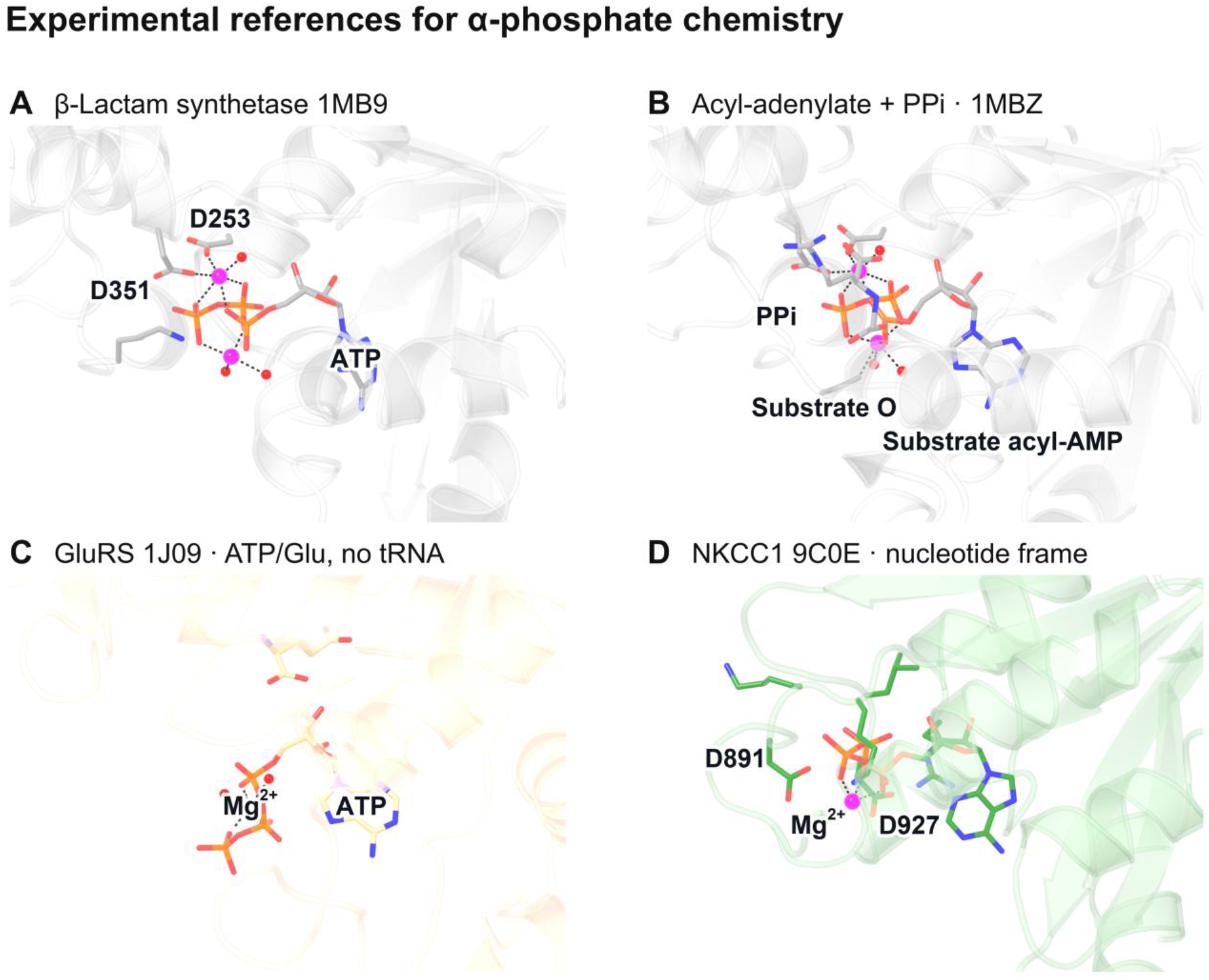
Experimental references for α-phosphate chemistry. (A) ATP-bound β-lactam synthetase 1MB9. (B) Its trapped substrate acyl-adenylate and PPi in 1MBZ (16). The covalently attached oxygen belongs to the small-molecule substrate. The OX2–Pα distance is 1.58 Å and the OX2–Pα–P(PPi) angle is 174°, describing product geometry. (C) Glutamyl-tRNA synthetase 1J09 with ATP and glutamate but no tRNA. This structure captures a nonproductive ATP-binding mode; tRNA binding switches ATP to a productive mode (17). (D) NKCC1 9C0E in the same nucleotide frame (8). Each structure retains its own modelled nucleotide and metal arrangement.

The SLC12 CTD contains two structurally related lobes. A global alignment can therefore place a single-domain partner on the lobe opposite the ATP site. After assigning partners to the ATP-binding N-lobe, lobe-level and nucleotide-frame comparisons locate ATP in the corresponding region of the fold (Figure 6). The eight-atom nucleotide fit to β-lactam synthetase has an RMSD of 0.34 Å. In that fixed frame, 108 of 123 N-lobe core Cα atoms lie within 5 Å of a β-lactam synthetase Cα atom, with a median nearest-neighbour distance of 3.0 Å. This metric describes spatial overlap after fitting the nucleotide; it does not define one-to-one residue equivalence.

**Figure 6.**
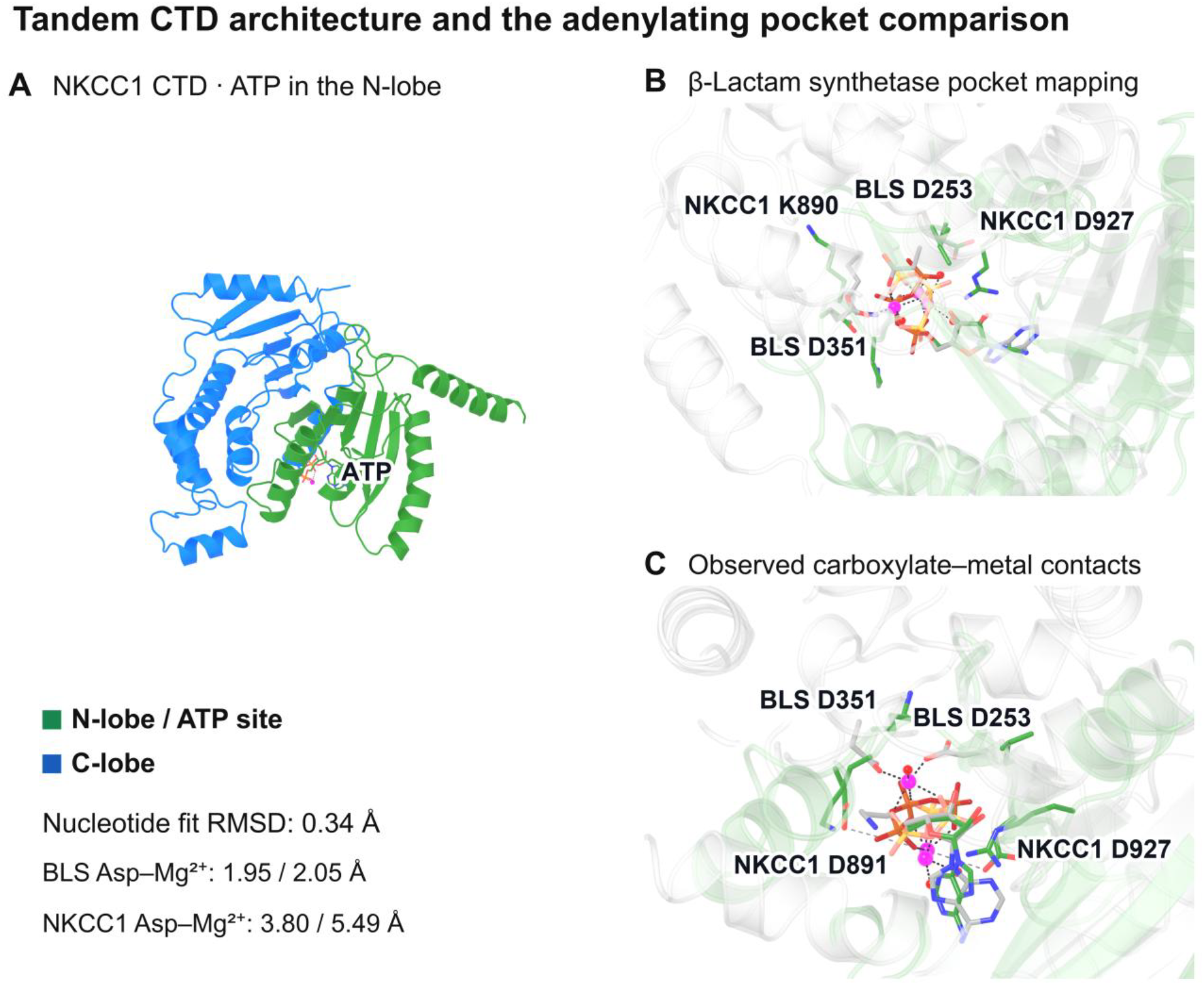
Tandem CTD architecture and the adenylating-pocket comparison. (A) NKCC1 9C0E CTD, with the ATP-binding N-lobe in green and the second lobe in blue. (B) β-Lactam synthetase 1MB9 (grey) in the NKCC1 nucleotide frame; the eight-atom ligand-fit RMSD is 0.34 Å. Its metal-binding Asp residues occupy the Lys890/Asp891 and Leu926/Asp927 pocket regions. This is a spatial comparison, and residue equivalence requires a separate alignment. (C) Observed metal contacts: β-lactam synthetase Asp253 OD2 and Asp351 OD1 lie 1.95 and 2.05 Å from Mg604, while the nearby NKCC1 carboxylates lie outside its first shell. Both structures retain their own modelled metals (8, 16). Lobe assignment precedes the comparison because a global DALI fit can place β-lactam synthetase on the second CTD lobe.

The β-lactam synthetase metal-binding Asp residues occupy regions beside NKCC1 Lys890/Asp891 and Leu926/Asp927 in the nucleotide frame (Figure 6). The nearby NKCC1 carboxylates remain outside the modelled Mg^2+^ first shell in 9C0E. A nearest-protein-carboxylate distance to Pα is not a direct comparison of the nucleophiles because β-lactam synthetase adenylates its substrate carboxylate. Glutamyl-tRNA synthetase supplies a different nucleotide–metal arrangement within the same broader fold (Figure 5). These comparisons locate the SLC12 nucleotide pocket relative to established α-phosphate chemistry and identify differences in its local coordination environment.

### UspG provides a protein adenylation precedent in a natural USP

Weber and Jung reported intrinsic autoadenylation and autophosphorylation of recombinant E. coli UspG. Chemical release followed by thin-layer chromatography identified covalently bound AMP (21). The acceptor residue remains unassigned. In the broader fold, bovine tryptophanyl-tRNA synthetase also forms protein-bound AMP and can regenerate ATP upon exposure to PPi (24). These observations make covalent AMP attachment a relevant biochemical endpoint for the SLC12 comparison.

Other USP studies measured different ATP-dependent signals. Bangera et al. reported ATPase activity for Salmonella YdaA using γ-^32^P-ATP thin-layer chromatography, assigning the released product to Pi (25). Xu et al. measured ATP remaining after incubation with E. coli UspE using Kinase-Glo (26). The UspE assay used 5 µM ATP and protein concentrations above ATP on a protomer basis, so ATP depletion at that endpoint does not quantify repeated turnover. MJ0577 showed extract-dependent ATP breakdown with an ADP assignment (22). The extract dependence leaves the contribution of purified MJ0577 unresolved.

These assays supply distinct levels of product information. UspG provides the protein–AMP precedent; the YdaA and UspE signals require coordinated nucleotide, phosphate and protein-adduct measurements to define their chemistry. Protein adenylation releases PPi while retaining AMP on the acceptor. Direct hydrolysis at Pα instead releases free AMP and PPi (Table S2). A γ-labelled ATP assay can report labelled Pi or PPi, depending on the cleaved bond, so the separation must resolve both products.

### Candidate acceptors require a change in geometry

The examined structures and predictions define the starting geometries for candidate protein acceptors (Figure 7). KCC2 9RNJ is an outward-facing T906A/T1007A construct at 3.86 Å resolution, with no modelled Mg^2+^ (27). Thr787 OG1 lies 4.1 Å from ATP Pα at an OG1–Pα–O3A angle of 123°. Across ten YdaA models, native Thr284 lies 4.2–4.6 Å from Pα at 120–123°. Explicit UspG ATP/Mg^2+^ predictions place Ser125 at 5.1–5.3 Å and 96–98° across 15 unique models and 25 nucleotide sites. These are ensemble ranges for YdaA and UspG; Table S1 reports the specified individual models.

**Figure 7.**
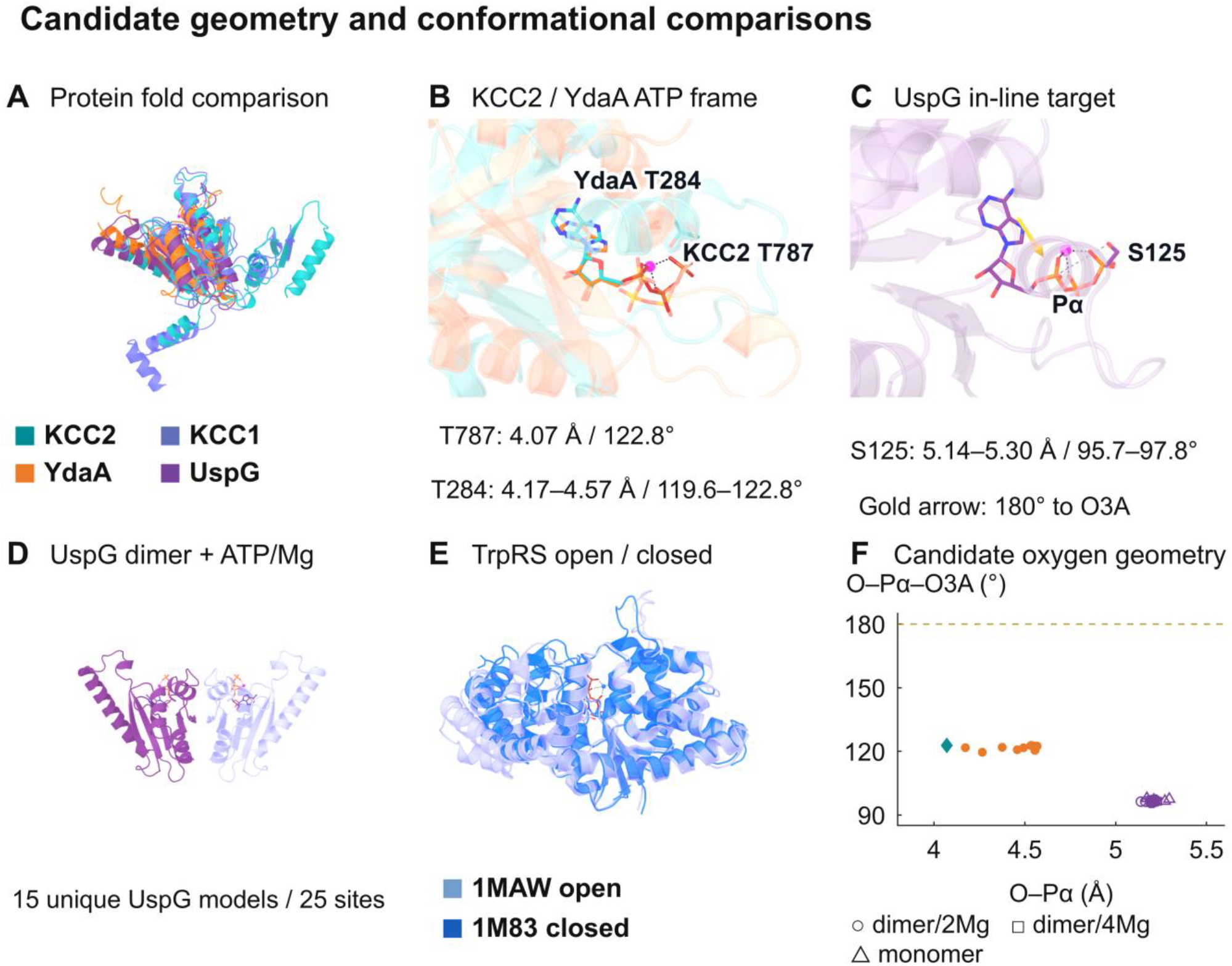
Candidate geometry and conformational comparisons. (A) Protein-only fold comparisons of KCC1/KCC2, YdaA and UspG. (B) Separate nucleotide-frame views of KCC2 Thr787 and native YdaA Thr284. KCC2 9RNJ is a 3.86-Å T906A/T1007A structure without modelled Mg^2+^ (27); YdaA ligand positions are predicted. (C) UspG Ser125 and a gold arrow along the 180° in-line direction at Pα, constructed without energy minimisation. (D) Natural UspG dimer with two ATP and two Mg^2+^, AlphaFold 3 model 0. (E) Bacterial TrpRS 1MAW open (light blue) and 1M83 closed (blue) ATP-bound states, aligned on catalytic-domain Cα atoms 1–200 (42). These structures illustrate conformational states during substrate activation. The bovine TrpRS protein-adenylation report is a separate biochemical observation (24). (F) Candidate oxygen–Pα distances and O–Pα–O3A angles. YdaA uses ten models; UspG uses 15 unique models and 25 sites, with conditions distinguished by marker. Models from one seed and paired dimer sites are correlated. The plotted residues are candidate acceptors, and the nucleotide-frame views do not establish residue homology.

With ATP held fixed, the candidate hydroxyls cannot reach an illustrative in-line point by side-chain rotation alone. A productive state would require backbone or nucleotide movement, or another acceptor. Changes in phosphorylation, transport state or protein association are possible triggers for testing. The additional ions in four-Mg^2+^ UspG predictions occupy variable positions and sometimes make implausibly short contacts. Those models therefore provide an uncertain description of extra-metal occupancy; Figure S3 shows the representative one-Mg-per-ATP pockets.

A conservation screen prioritises hydroxyl-bearing residues for biochemical comparison (Table S1). In the principal query-centred pools, Ser/Thr retention is 93.9% at UspG S125, 94.7% at S113, 97.4% at YdaA T284, 90.7% at T270 and 94.7% at S277. UspG S125 corresponds to YnaF S127, which retains Ser/Thr in 84.0% of its pool. Wild-type YnaF bound ATP and showed no detectable ATPase activity under the reported assay conditions (25). YnaF therefore provides a conserved-site comparison for experiments that measure the same ATP-derived products under matched conditions. The UspG S125–YdaA T284 pairing is not an established sequence correspondence; the SI gives the residue mappings and additional candidates.

## Discussion

Both SLC12 CTD lobes score between USPs and EC-annotated domains in the fold comparison, and the N-lobe places its nucleotide in a region used for adenylation by catalytic members of the broader fold. Our analysis quantifies this relationship and identifies paralog-specific changes around the phosphate chain. NKCC1 retains the D891/D927/R801/K889 arrangement, while KCC2 replaces or loses several corresponding groups. These differences parallel the distinct responses of NKCCs and KCCs to WNK/SPAK/OSR1 phosphorylation and may support different regulatory functions at their nucleotide sites (4).

The CTD consists of two lobes of the same fold. Each SLC12 lobe groups with the corresponding lobe of the archaeal CCC domain in the DALI comparison, and bacterial SLC12 homologues contribute lobes to both lobe clades of the tree. The two lobes have been separate since before the divergence of the sampled archaeal, bacterial and eukaryotic CCCs, and ATP binds the N-lobe in the examined structures.

USP biology connects nucleotide recognition with transport regulation and protein interactions. TeaD regulates the ectoine transporter TeaABC (23). The USP domain of Staphylococcus aureus KdpD binds c-di-AMP and regulates expression of the Kdp potassium transporter (28). More recently, a Klebsiella quasipneumoniae UspG homologue was shown to bind c-di-GMP and promote association with the transcriptional regulator RamR (29). These examples illustrate how USP nucleotide recognition can influence associated proteins. Their ligand specificities belong to the proteins tested and provide functional context for investigating SLC12 regulation.

The UspG protein–AMP result motivates a specific chemical hypothesis for the transporter site. The geometrical requirement is an acceptor positioned near Pα and approximately in line with the leaving PPi group. Glutamyl-tRNA synthetase demonstrates how binding of a partner, tRNA, can switch ATP into a productive binding mode (17). This supplies a direct precedent for conditional nucleotide geometry in the broader fold. The candidate geometries in SLC12 and USP models identify the rearrangements to test; the structures do not determine the trigger or energetics of such a state.

For UspG, separate S125A and S113A comparisons would test the leading hydroxyl candidates. YdaA T284A, T270A and S277A would become informative after a wild-type covalent protein–AMP product is established. Site-resolved adduct identification, together with ATP-binding, folding and oligomerisation measurements, would distinguish an acceptor substitution from a change in pocket organisation. The SI sets out product measurements, adduct-enrichment options, linkage-sensitive detection and ATP-analogue controls (Tables S1 and S2). Parallel assays of a soluble CTD and the full-length transporter would test the contribution of membrane-protein context to any detected chemistry.

Protein adenylation at an SLC12 nucleotide site remains a testable proposal. A direct test should identify the covalent protein–AMP adduct and its acceptor, measure PPi and resolve the soluble ATP, ADP, AMP and Pi products. Quantitative time courses would measure adduct lifetime and ATP consumption per protomer, providing tests of transient attachment and repeated turnover. These measurements would determine how the structural relationship to USPs and adenylating enzymes translates into nucleotide-site function.

## Methods

### Family alignment

We retrieved 26,832 UniProt sequences cross-referenced to InterPro IPR004842 and selected 16,192 CTD-containing sequences using a length threshold of ≥700 amino acids and a C-terminal DGG-motif filter (18, 30). The inputs were CTD-containing protein sequences, not excised CTDs. These filters restrict the analysis to long proteins retaining the selected motif. We computed the multiple-sequence alignment with pyFAMSA 0.7.0, using the medoid-tree heuristic and 12 threads (31, 32). The alignment contains 42,533 columns. KCC1 D804, NKCC1 D891 and NKCC1 D927 map to columns 25651, 25532 and 26926, respectively. Per-column residue counts and conditional probabilities were computed in Python. The reported overall Asp frequencies use non-gap entries at each column. Clade assignment used keyword parsing of UniProt OS= fields; Figure S1 includes alignment gaps in each clade denominator. Table 1 defines chemical classes for pairs of known amino acids and does not report inferred class frequencies.

### AlphaFold 3 modelling

We used the AlphaFold Server to predict dimeric human KCC2b (1,116 amino acids; UniProt Q9H2X9-2) and, for comparison, KCC2a (1,139 amino acids; Q9H2X9-1) (33). All KCC2 inputs also contained human creatine kinase B (381 amino acids), as two copies in most jobs. The models used for the reported KCC2 geometry contained ATP and Mg^2+^, with phosphothreonine 906 and phosphoserine 1045 in phosphorylated KCC2b (929 and 1068 in KCC2a). A later set modelled phosphothreonine 906 and 1007 singly and together, with ATP and without Mg^2+^. Each archived job returned five ranked models from one seed. The original comparisons used the highest-ranked model; seed values are recorded in the saved job requests. Residue numbering follows KCC2b throughout. KCC2a positions are offset by +23 residues, so KCC2b position X corresponds to KCC2a X+23. The inhibitory KCC2b phosphosites are T906 and T1007. Pocket distances were extracted from mmCIF coordinates with a custom script.

### Structural survey

We retrieved 883,981 UniProtKB sequences carrying InterPro IPR014729 and reduced them with hierarchical CD-HIT clustering at 90%, 70% and 50% identity to 56,437 representatives (34). We retrieved SLC12 sequences separately through Pfam PF03522, yielding 19,939 sequences and 935 representatives at 50% identity. The two retrieval sets contained 57,372 representatives in total. AlphaFold Protein Structure Database models were available for 43,988 representatives (76.7%), and 42,845 produced DALI alignments (35). This annotation-derived survey defines the scope of the fold comparison.

### DALI scores and grouping

We ran DaliLite v5 locally (36). MJ0577 versus TeaD gave Z = 18.5 in both directions, as on the DALI web server (37). An absent alignment contributes Z = 0, with an RMSD of 20 Å in the clustering matrix. The score is the mean Z to MJ0577, TeaD and the higher-scoring archaeal CCC lobe divided by the mean Z to the β-lactam synthetase and glutamyl-tRNA synthetase domains. Of the 46,367 domains from the survey, 45,378 have a nonzero denominator and a defined score. Each protein carries one group label: SLC12 (Pfam PF03522 retrieval), USP (Pfam PF00582), EC-annotated (an EC number and no PF00582) or other. Group comparisons used common-language effect sizes with ties counted at half weight and 95% confidence intervals from 1,000 bootstrap resamples. The seven scoring rules were the mean ratio, the ratio with one added to the denominator, the mean ratio restricted to domains aligned to both enzyme references or to all references, the ratio of the highest Z values, the difference of means and the difference of the highest Z values. The k-means analysis used column-standardised Z and RMSD profiles; k was selected by adjusted Rand agreement with the annotation groups.

### Fold domains

For each of the 42,845 models, we defined the fold domain from the DALI equivalences of the whole-protein survey to MJ0577, or, where that Z was below 2, to the highest-scoring other reference within its catalytic core. In whole-protein alignments, the references often matched different SLC12 lobes (Figure S4). The domain is the largest cluster of aligned blocks, padded by five residues, with unaligned internal segments longer than 40 residues removed (42,821 domains; median 149 residues). Second lobes of SLC12 and tandem USP proteins were located with Foldseek (38) using experimental lobe templates, and SLC12 lobes were assigned as N- or C-lobes from Foldseek scores against the 9C0E and 7AIP lobes. After removal of domains shorter than 50 residues, 46,380 domains remained, including 13 experimental domains used as landmarks. Each domain was compared with single-domain references: MJ0577 (1MJH chain A), TeaD (3HGM A), the archaeal CCC N- and C-lobes (3G40 A 492–627 and 628–760), β-lactam synthetase (1MB9 A 210–507) and glutamyl-tRNA synthetase (1J09 A 1–305). The last two follow SCOPe domain boundaries (15). Median Z values for each reference are given in Table S3.

### Lobe assignment and nucleotide fitting

We assigned structural partners to a CTD lobe by counting DALI-equivalent residues on either side of NKCC1 residue 980. Partners assigned to the ATP-binding N-lobe were used for its protein comparison. Nucleotide-frame fits used C1′, C2′, C3′, C4′, O4′, C5′, O5′ and Pα. The 0.34-Å RMSD reports the fitted nucleotide atoms. The separate N-lobe overlap statistic is a nearest-neighbour Cα distance in that fixed frame; it is not a residue-paired protein-fit RMSD. We screened first-shell metal contacts within 2.6 Å after removal of riding hydrogens and classified ions by their phosphate contacts following Dudas et al. (39). The N-lobe core comprises the 123 NKCC1 residues between 786 and 924 that are DALI-equivalent to MJ0577 or YnaF.

### Structural trees

Domains were clustered with Foldseek (coverage 0.7, identity 0.2). Representatives were chosen from distinct clusters within each family, with both lobes of 200 SLC12 proteins and 150 tandem USPs; 1,977 passed the length filter. Exhaustive all-against-all Foldseek alignments (3Di+AA) were converted to 1 − LDDT distances, with 1 − mean TM-score and −ln(sequence identity) as alternatives, and trees were built with FastME (balanced minimum-evolution taxon addition followed by balanced NNI) (40, 41). For display, trees were rooted on the branch that best separates the nucleotidylyl-transferase superfamily from the remaining domains. Each surveyed domain was placed on its nearest representative by Foldseek bit score (Figure S5, Table S4).

### Additional structural comparisons

We used experimental KCC2 9RNJ, KCC1 7AIP and YdaA 4R2J, the AlphaFold DB model of E. coli UspG P39177, and the archived YdaA ATP jobs with one or two Mg^2+^ ions (5, 25, 27). Native YdaA numbering is restored by adding one to the original input positions. The UspG predictions used the native 142-residue sequence with dimer/2ATP/2Mg^2+^, monomer/ATP/Mg^2+^ and dimer/2ATP/4Mg^2+^ inputs at seeds 10501, 10502 and 10503. Each input returned five ranked models. Deduplication by atom identity and position retained 15 UspG models and 25 sites; the two YdaA jobs provided ten models. Distances are acceptor-to-Pα and angles are acceptor–Pα–O3A. The 3.2-Å, 180° in-line point is an illustrative geometric construction. Waters and reaction paths were not modelled. The SI identifies the individual models used in Table S1 and describes confidence, metal-contact and conservation analyses. Bacterial TrpRS 1MAW and 1M83 were aligned on catalytic-domain Cα atoms 1–200 for the conformational comparison in Figure 7 (42).

### USP conservation

We excluded query copies and duplicate aligned rows from the archived AlphaFold search alignments. Principal pools required ≥40% identity and ≥80% query coverage, yielding 415 UspG, 787 YdaA and 566 YnaF sequences. Henikoff-weighted residue frequencies, broader-pool and redundancy checks, native numbering and per-site occupancy are detailed in the SI. These query-centred pools contain homologues and are not curated orthologue sets.

## Data and code availability

The analysis records comprise the input sequence lists, filtered alignments and anchor mappings, structural-source manifests, AlphaFold job requests and coordinates, residue-frequency tables and analysis scripts. The USP conservation file inventory is given in the Supplementary Information.

## References

1. Rivera C, Voipio J, Payne JA, Ruusuvuori E, Lahtinen H, Lamsa K, et al. (1999). The K+/Cl™ co-transporter KCC2 renders GABA hyperpolarizing during neuronal maturation. Nature 397:251–255. 10.1038/16697

2. Kaila K, Price TJ, Payne JA, Puskarjov M, Voipio J (2014). Cation-chloride cotransporters in neuronal development, plasticity and disease. Nat Rev Neurosci 15:637–654. 10.1038/nrn3819

3. Medina I, Friedel P, Rivera C, Kahle KT, Kourdougli N, Uvarov P, et al. (2014). Current view on the functional regulation of the neuronal K+-Cl™ cotransporter KCC2. Front Cell Neurosci 8:27. 10.3389/fncel.2014.00027

4. Alessi DR, Zhang J, Khanna A, Hochdörfer T, Shang Y, Kahle KT (2014). The WNK-SPAK/OSR1 pathway: Master regulator of cation-chloride cotransporters. Sci Signal 7:re3. 10.1126/scisignal.2005365

5. Chi G, Ebenhoch R, Man H, Tang H, Tremblay LE, Reggiano G, et al. (2021). Phospho-regulation, nucleotide binding and ion access control in potassium-chloride cotransporters. EMBO J 40:e107294. 10.15252/embj.2020107294

6. Fan M, Zhang J, Lee CL, Zhang J, Feng L (2023). Structure and thiazide inhibition mechanism of the human Na–Cl cotransporter. Nature 614:788–793. 10.1038/s41586-023-05718-0

7. Zhao Y, Schubert H, Blakely A, Forbush B, Smith MD, Rinehart J, et al. (2024). Structural bases for Na+-Cl™ cotransporter inhibition by thiazide diuretic drugs and activation by kinases. Nat Commun 15:7006. 10.1038/s41467-024-51381-y

8. Zhao Y, Vidossich P, Forbush B, Ma J, Rinehart J, De Vivo M, et al. (2025). Structural basis for human NKCC1 inhibition by loop diuretic drugs. EMBO J 44:1540–1562. 10.1038/s44318-025-00368-6

9. Warmuth S, Zimmermann I, Dutzler R (2009). X-ray Structure of the C-Terminal Domain of a Prokaryotic Cation-Chloride Cotransporter. Structure 17:538–546. 10.1016/j.str.2009.02.009

10. Zimanyi CM, Guo M, Mahmood A, Hendrickson WA, Hirsh D, Cheung J (2020). Structure of the Regulatory Cytosolic Domain of a Eukaryotic Potassium-Chloride Cotransporter. Structure 28:1051–1060.e4. 10.1016/j.str.2020.06.009

11. Chi X, Li X, Chen Y, Zhang Y, Su Q, Zhou Q (2021). Cryo-EM structures of the full-length human KCC2 and KCC3 cation-chloride cotransporters. Cell Res 31:482–484. 10.1038/s41422-020-00437-x

12. Xie Y, Chang S, Zhao C, Wang F, Liu S, Wang J, et al. (2020). Structures and an activation mechanism of human potassium-chloride cotransporters. Sci Adv 6:eabc5883. 10.1126/sciadv.abc5883

13. Chew TA, Orlando BJ, Zhang J, Latorraca NR, Wang A, Hollingsworth SA, et al. (2019). Structure and mechanism of the cation–chloride cotransporter NKCC1. Nature 572:488–492. 10.1038/s41586-019-1438-2

14. Aravind L, Anantharaman V, Koonin EV (2002). Monophyly of class I aminoacyl tRNA synthetase, USPA, ETFP, photolyase, and PP-ATPase nucleotide-binding domains: implications for protein evolution in the RNA world. Proteins 48:1–14. 10.1002/prot.10064

15. SCOPe. SCOPe structural classification hierarchy: adenine nucleotide alpha hydrolase-like fold (c.26) (version 2.08; accessed 2026-10-12). https://scop.berkeley.edu/

16. Miller MT, Bachmann BO, Townsend CA, Rosenzweig AC (2002). The catalytic cycle of β-lactam synthetase observed by x-ray crystallographic snapshots. Proc Natl Acad Sci USA 99:14752–14757. 10.1073/pnas.232361199

17. Sekine S, Nureki O, Dubois DY, Bernier S, Chênevert R, Lapointe J, et al. (2003). ATP binding by glutamyl-tRNA synthetase is switched to the productive mode by tRNA binding. EMBO J 22:676–688. 10.1093/emboj/cdg053

18. Blum M, Andreeva A, Florentino LC, Chuguransky SR, Grego T, Hobbs E, et al. (2025). InterPro: the protein sequence classification resource in 2025. Nucleic Acids Res 53:D444–D456. 10.1093/nar/gkae1082

19. Dessailly BH, Redfern OC, Cuff AL, Orengo CA (2010). Detailed Analysis of Function Divergence in a Large and Diverse Domain Superfamily: Toward a Refined Protocol of Function Classification. Structure 18:1522–1535. 10.1016/j.str.2010.08.017

20. Yin J, Waman VP, Sen N, Firdaus-Raih M, Lam SD, Orengo C (2025). Understanding the structural and functional diversity of ATP-PPases using protein domains and functional families in the CATH database. Structure 33:613–631.e6. 10.1016/j.str.2024.12.016

21. Weber A, Jung K (2006). Biochemical Properties of UspG, a Universal Stress Protein of Escherichia coli. Biochemistry 45:1620–1628. 10.1021/bi051301u

22. Zarembinski TI, Hung LW, Mueller-Dieckmann HJ, Kim KK, Yokota H, Kim R, et al. (1998). Structure-based assignment of the biochemical function of a hypothetical protein: A test case of structural genomics. Proc Natl Acad Sci USA 95:15189–15193. 10.1073/pnas.95.26.15189

23. Schweikhard ES, Kuhlmann SI, Kunte HJ, Grammann K, Ziegler CM (2010). Structure and Function of the Universal Stress Protein TeaD and Its Role in Regulating the Ectoine Transporter TeaABC of Halomonas elongata DSM 2581T. Biochemistry 49:2194–2204. 10.1021/bi9017522

24. Merkulova TI, Kovaleva GK, Kiselev LL (1987). Autoadenylation of tryptophanyl-tRNA synthetase. Mol Biol (Mosk) 21:769–776. PMID: 3657776. https://pubmed.ncbi.nlm.nih.gov/3657776/

25. Bangera M, Panigrahi R, Sagurthi SR, Savithri HS, Murthy MRN (2015). Structural and functional analysis of two universal stress proteins YdaA and YnaF from Salmonella typhimurium: possible roles in microbial stress tolerance. J Struct Biol 189:238–250. 10.1016/j.jsb.2015.01.003

26. Xu Y, Guo J, Jin X, Kim JS, Ji Y, Fan S, et al. (2016). Crystal structure and functional implications of the tandem-type universal stress protein UspE from Escherichia coli. BMC Struct Biol 16:3. 10.1186/s12900-016-0053-9

27. Matsuoka R, Oswald C, Jazayeri A, Duerr KL (2026). Cryo-EM structure of the human potassium chloride cotransporter T906A/T1007A phospho-knockout mutants KCC2b bound ATP in LMNG (outward-facing state, dimer) [PDB 9RNJ, version 1.0]. 10.2210/pdb9RNJ/pdb

28. Moscoso JA, Schramke H, Zhang Y, Tosi T, Dehbi A, Jung K, et al. (2016). Binding of Cyclic Di-AMP to the Staphylococcus aureus Sensor Kinase KdpD Occurs via the Universal Stress Protein Domain and Downregulates the Expression of the Kdp Potassium Transporter. J Bacteriol 198:98–110. 10.1128/JB.00480-15

29. Liu X, Wang M, Fu Y, Zhu X, Du M, Wen Y, et al. (2026). Cyclic di-GMP directly reprograms the multidrug resistance machinery via UspG-mediated sequestration of RamR in Klebsiella. PLoS Pathog 22:e1014537. 10.1371/journal.ppat.1014537

30. The UniProt Consortium (2025). UniProt: the Universal Protein Knowledgebase in 2025. Nucleic Acids Res 53:D609–D617. 10.1093/nar/gkae1010

31. Larralde M (2026). PyFAMSA: Cython bindings and Python interface to FAMSA (version 0.7.0) [software]. https://pypi.org/project/pyfamsa/0.7.0/

32. Gudyś A, Zielezinski A, Notredame C, Deorowicz S (2026). Fast and accurate multiple-protein-sequence alignment at scale with FAMSA2. Nat Biotechnol Advance online publication. 10.1038/s41587-026-03095-3

33. Abramson J, Adler J, Dunger J, Evans R, Green T, Pritzel A, et al. (2024). Accurate structure prediction of biomolecular interactions with AlphaFold 3. Nature 630:493–500. 10.1038/s41586-024-07487-w

34. Li W, Godzik A (2006). Cd-hit: a fast program for clustering and comparing large sets of protein or nucleotide sequences. Bioinformatics 22:1658–1659. 10.1093/bioinformatics/btl158

35. Varadi M, Bertoni D, Magana P, Paramval U, Pidruchna I, Radhakrishnan M, et al. (2024). AlphaFold Protein Structure Database in 2024: providing structure coverage for over 214 million protein sequences. Nucleic Acids Res 52:D368–D375. 10.1093/nar/gkad1011

36. Holm L (2019). Benchmarking fold detection by DaliLite v.5. Bioinformatics 35:5326–5327. 10.1093/bioinformatics/btz536

37. Holm L (2022). Dali server: structural unification of protein families. Nucleic Acids Res 50:W210–W215. 10.1093/nar/gkac387

38. van Kempen M, Kim SS, Tumescheit C, Mirdita M, Lee J, Gilchrist CLM, et al. (2024). Fast and accurate protein structure search with Foldseek. Nat Biotechnol 42:243–246. 10.1038/s41587-023-01773-0

39. Dudas B, Berta D, Jambrina P, Buigues PJ, Suardiaz R, Gómez-Coca S, et al. (2026). Nature Counts to Three: Universal Mg-Pinch Motif Polarizes the Cleaved Bond in NTP-Processing Enzymes. J Am Chem Soc 148:30494–30511. 10.1021/jacs.6c10146

40. Lefort V, Desper R, Gascuel O (2015). FastME 2.0: A Comprehensive, Accurate, and Fast Distance-Based Phylogeny Inference Program. Mol Biol Evol 32:2798–2800. 10.1093/molbev/msv150

41. Moi D, Bernard C, Steinegger M, Nevers Y, Langleib M, Dessimoz C (2025). Structural phylogenetics unravels the evolutionary diversification of communication systems in gram-positive bacteria and their viruses. Nat Struct Mol Biol 32:2492–2502. 10.1038/s41594-025-01649-8

42. Retailleau P, Huang X, Yin Y, Hu M, Weinreb V, Vachette P, et al. (2003). Interconversion of ATP Binding and Conformational Free Energies by Tryptophanyl-tRNA Synthetase: Structures of ATP Bound to Open and Closed, Pre-Transition-state Conformations. J Mol Biol 325:39–63. 10.1016/S0022-2836(02)01156-7

