## Supplementary Information for "Structural relationships between the SLC12 cotransporter C-terminal domain, universal stress proteins and adenylating enzymes"

We screened potential protein nucleophiles in universal stress proteins (USPs) to prioritise residues for biochemical tests of AMP attachment. UspG S125 and S113, and YdaA T284, T270 and S277 retain hydroxyl groups in their respective sequence pools. Their positions in the examined coordinates require a change in protein or nucleotide geometry before these hydroxyls can attack ATP Pα in line with the departing group. Table S1 presents these candidates with their conservation and geometry. Published experiments identify protein-bound AMP in recombinant E. coli UspG; the YdaA and UspE studies report ATP-dependent signals with less complete product identification (1, 2, 3).

### Sequence conservation methods

We reused the unpaired A3M search alignments from the archived AlphaFold 3 jobs for E. coli K-12 UspG (P39177), Salmonella Typhimurium YdaA (Q8ZP84) and YnaF (P67091). We removed lowercase insertions and period characters to recover query-aligned columns, then excluded the query, every exact query copy and duplicate aligned rows. The principal pools retained sequences with ≥40% exact identity to the query and ≥80% query coverage, yielding 415 UspG, 787 YdaA and 566 YnaF aligned sequences. Identity was the number of exact matches divided by the number of positions containing known aligned amino acids. Coverage was the fraction of query positions occupied by known amino acids. Gaps and ambiguous residues contributed neither exact matches nor occupied positions.

We calculated position-based Henikoff weights across each retained alignment (4). We normalised site frequencies to the sum of weights for known amino acids at that position. Table S1 reports exact residue retention, Ser/Thr retention for hydroxyl candidates, and the unweighted number of known residues over the retained pool size. These search-hit pools provide descriptive conservation estimates. They contain uncurated homologues and related sequences, and the analysis includes no phylogenetic correction or test of selection.

The sensitivity analyses lowered the identity threshold to 25% at the same coverage, yielding 11,025 UspG, 3,675 YdaA and 10,311 YnaF sequences. Within the principal pools, we also used inverse 80%-identity-neighbourhood weights and unweighted greedy 80%-identity representatives. The latter retained 126, 273 and 290 representatives, respectively. The source dataset retains the site frequencies, redundancy checks and annotation-restricted comparisons.

### Numbering coordinates and geometry

Numbering follows the native UniProt sequences, including the initiator methionine. The YdaA and YnaF AlphaFold inputs omit this residue, so their model positions require a +1 offset; UspG requires no offset. Table S1 uses the UspG dimer/two-Mg model 0, YdaA one-Mg model 0 and experimental YnaF 4R2L chain A (2). The UspG and YdaA nucleotide positions come from predictions.

The archived UspG AlphaFold 3 calculations used the native 142-residue E. coli sequence P39177 (5). Inputs comprised a dimer with two ATP and two $Mg^{2+}$ ions (seed 10501), a monomer with ATP and $Mg^{2+}$ (seed 10502), and a dimer with two ATP and four $Mg^{2+}$ ions (seed 10503). Each job returned five ranked models. The two archived YdaA jobs, with one or two $Mg^{2+}$ ions, also returned five models each. Coordinate deduplication by atom identity and position retained 15 UspG models and 25 sites for the geometric summaries in the main text; the YdaA comparison used ten models. Models from a single seed and sites within one dimer share a prediction context and provide correlated observations.

We measured the distance from each listed potential nucleophilic atom to ATP Pα and the atom–Pα–O3A angle. O3A is the oxygen of the α–β phosphate bridge and marks the leaving-group direction for attack at Pα. For His and carboxylates, Table S1 reports the atom closer to Pα; the source data retain both alternatives. Protein comparisons use the corresponding ATP-binding lobes. Nucleotide superpositions use eight atoms: C1′, C2′, C3′, C4′, O4′, C5′, O5′ and Pα. The reported distances and angles describe individual static coordinates; their decimal places do not imply experimental or predictive accuracy at that precision.

The geometric screen evaluates potential protein acceptors. Water-mediated ATP hydrolysis requires a separate assessment of water occupancy and catalytic geometry. These models contain no constructed catalytic water or predicted reactive state. The main-text in-line point is a geometric construction at 3.2 Å from Pα and 180° to the leaving-group direction, without energy minimisation. We measured ATP–protein heavy-atom distances and separately screened $Mg^{2+}$ contacts to O, N or S within 2.6 Å. Variable additional $Mg^{2+}$ positions, including implausibly short contacts in some four-Mg predictions, remain model features with uncertain physical interpretation.

UspG S125 and YnaF S127 correspond in both the protein structural comparison and an independent sequence alignment. We treat UspG S125 and YdaA T284 as separate pocket candidates: the sequence alignment places UspG S125 against YdaA N283. Local gaps affect the S113/T270 mapping. The YdaA crystal structure lacks density for residues 272–278, including S277 (2). The S277 geometry in Table S1 comes from a predicted loop with low confidence (pLDDT ≈47).

### Published biochemical evidence

Weber and Jung reported autoadenylation and autophosphorylation of recombinant E. coli UspG, with chemical release and thin-layer chromatography identifying covalently bound AMP (1). The acceptor residue remains unassigned. Bangera et al. reported ATPase activity for Salmonella YdaA using γ-$^{32}$P-ATP, whereas wild-type YnaF bound ATP and showed no detectable ATPase activity under their assay conditions (2). YnaF therefore provides an ATP-binding comparison with a defined assay history.

Xu et al. measured ATP remaining after incubation with E. coli UspE using Kinase-Glo (3). Their 20-minute endpoint used 5 µM ATP and 0.4–3.2 mg/ml protein. These conditions permit stoichiometric ATP loss and do not quantify repeated catalytic turnover. Identification of the adenine-nucleotide and inorganic-phosphate products would resolve the reaction chemistry behind the YdaA and UspE signals.

### Proposed residue comparisons

For purified E. coli K-12 UspG, we propose testing S125A and S113A separately against wild type. S125 lies relatively near ATP and contacts a γ-phosphate oxygen in the model. S113 retains a hydroxyl strongly across the principal sequence pool, but lies buried and farther from Pα. Each comparison should measure protein-bound AMP alongside ATP binding, structural integrity and oligomeric state. A reduction in labelling could result from loss of the acceptor, altered nucleotide recognition or a conformational change. T120A is a secondary comparison because hydroxyl retention falls sharply in the broader pool. S128A provides a lower-conservation hydroxyl comparison; its structural and biochemical effects would require the same controls.

For YdaA, the first proposed endpoint is a covalent protein–AMP product in wild type. Detection of this product would support individual tests of T284A, T270A and S277A. These positions also map to T284, T270 and S277 in reviewed E. coli K-12 UspE (P0AAC0), allowing a parallel comparison in that homologue. The Salmonella structural results and E. coli biochemical results describe separate proteins. A comparison with wild-type YnaF and its S127A variant could test the structurally and sequence-supported S125/S127 correspondence; YnaF activity would need to be measured under the same conditions as UspG.

UspG H121 and YdaA H169 are secondary candidates. Their positions support phosphate contacts, and site-resolved adduct identification could test whether either acts as an AMP acceptor. YdaA H225, H244 and C294 are retained but distant from Pα and partly buried, giving them lower priority in the current coordinates. UspG D10 and YdaA E286 provide carboxylate groups with possible binding, coordination or distinct covalent-linkage roles. The strongly retained YdaA T128 lies in the other domain, about 21 Å from the modelled C-terminal ATP site.

### Proposed product identification

Assigning an acceptor requires identification of a covalent protein–AMP adduct and localisation of the modified residue. We propose complementary intact-protein and peptide mass spectrometry after removal of noncovalently bound nucleotide. Protein labelling from α-labelled ATP could provide an additional readout, with parallel analysis for phosphorylation and retained ATP. An intact mass shift consistent with AMP would support adduct formation; peptide-level evidence would locate its attachment site. Mutant effects should be interpreted together with nucleotide binding and structural measurements.

An engineered hinT affinity reagent could enrich low-occupancy AMPylated proteins for subsequent analysis, following Gonzalez et al. (6). A validated protein–AMP positive standard would test capture efficiency for the linkage and substrate under study. Enrichment would supplement direct adduct identification. Zhang et al. captured a His–AMP phosphoramidate linkage in an S261A mutant of the bacterial effector LnaB (7), providing a chemical precedent for the histidine candidates and a reason to preserve potentially labile linkages during sample preparation. A protein-bound AMP species could represent a catalytic intermediate or a persistent modification; time courses would measure adduct lifetime, while kinetic and functional tests would assess its role. These studies guide the detection and handling of protein–AMP linkages.

For protein adenylation from ATP, the immediate products are protein–AMP and PPi. Free AMP can arise from deAMPylation, hydrolysis of an intermediate or direct ATP hydrolysis at Pα. We propose measuring ATP, ADP and AMP separately from Pi, PPi and the protein adduct, with time courses to detect subsequent PPi hydrolysis or adduct loss. Table S2 summarises the primary products that distinguish these processes.

#### Table S2 Primary products for distinguishing ATP binding and chemistry

This schematic comparison defines prospective biochemical readouts. It presents possible reactions, with protonation states omitted.

| Process | Nucleophile and attacked phosphate | Other primary product(s) | Protein-associated product |
|---|---|---|---|
| ATP binding | No covalent attack | None | Noncovalent protein·ATP |
| ATP hydrolysis to ADP | Water attacks Pγ | ADP + Pi | Unmodified protein |
| Protein phosphorylation | Protein nucleophile attacks Pγ | ADP | Covalent protein–phosphate |
| ATP hydrolysis to AMP | Water attacks Pα | AMP + PPi | Unmodified protein |
| Protein adenylation | Protein nucleophile attacks Pα | PPi | Covalent protein–AMP |
| Small-molecule adenylation | Small-molecule nucleophile attacks Pα | Substrate–AMP + PPi | No covalent protein adduct required |

A substrate–AMP intermediate can remain bound to the enzyme without a covalent protein–AMP linkage. The substrate and protein species should therefore be distinguished during product analysis.

Release of the γ label from γ-$^{32}$P-ATP can produce labelled Pi after β–γ cleavage or labelled PPi after α–β cleavage. Chromatography with both standards would distinguish these products. ATP depletion alone identifies neither the cleaved bond nor the acceptor.

### Proposed assay controls and ATP analogue comparisons

The proposed time courses should establish adduct stoichiometry per protomer and distinguish a single modification event from repeated ATP turnover. Protein concentrations below the ATP concentration would help assess sustained consumption. ATP-spike recovery and addition of protein after reaction quenching would test interference with an ATP-detection assay. Activity should track the target protein across purification, and assays should include matched buffer, no-protein and protein-integrity controls. These measurements would connect the product signal to the purified protein and its binding-competent state.

ATP analogues should be chosen according to the phosphate bridge under study. AMP-PNP replaces the β–γ bridging oxygen with NH, leaving the α–β bridge intact: He et al. detected AMP production from AMP-PNP by actin-activated Ceg14 (8). Comparisons of analogues modified at the α–β and β–γ bridges would therefore be useful alongside ATP, with binding assessed independently. Changes in analogue activity can reflect altered recognition as well as a blocked chemical step. The Ceg14 result provides a precedent for interpreting these chemical probes.

### Guided structure predictions

We also tested restraint-guided structure predictions with Boltz-2 as a way to reach an in-line geometry at UspG (9, 10). Across 60 models with distance or angle restraints on S125, S113 or T120, 20 control models and 12 models of a further pose pilot, no model satisfied both the geometric window (3.0–3.5 Å, 160–180°) and a screen for Mg–oxygen contacts shorter than 1.8 Å. One pilot model reached 3.48 Å and 166° at S125 but places a $Mg^{2+}$ ion 0.19 Å from an ATP oxygen. Among the unguided controls, 0 of 10 models have a Mg–oxygen contact below 1.8 Å, against 5 of 10 for the geometry-only arm. These runs describe the behaviour of the prediction protocol under restraints and do not measure UspG activity.

### Associated data

The USP conservation dataset comprises candidate_shortlist.csv, candidate_summary.csv, all_position_conservation.csv, redundancy_robustness.csv, reviewed_reference_mapping.csv, filtered alignments and input manifests. The structural source archive contains the coordinates and original AlphaFold search alignments. These records support the residue frequencies, model identities, numbering offsets and geometric comparisons described here.

### Table S1 Conservation and geometry of candidate USP nucleophiles

Exact and Ser/Thr frequencies are Henikoff-weighted percentages. Broad retention uses the ≥25% identity pool; all other conservation columns use the ≥40% pool.

| Protein site | Known / N | Exact % | Ser/Thr % | Broad retention % | Candidate atom | Pα distance Å | Angle ° | Interpretation |
|---|---|---|---|---|---|---|---|---|
| UspG S125 | 398/415 | 92.6 | 93.9 | 83.2 | OG | 5.14 | 96.3 | First hydroxyl candidate; contacts a γ-phosphate oxygen. |
| UspG S113 | 415/415 | 88.8 | 94.7 | 74.6 | OG | 8.43 | 59.1 | Conserved hydroxyl; buried in the model. |
| UspG T120 | 404/415 | 77.1 | 83.3 | 14.2 | OG1 | 10.40 | 24.1 | Secondary hydroxyl candidate; lineage dependent. |
| UspG H121 | 403/415 | 46.2 | — | 11.7 | NE2 | 5.87 | 19.8 | Secondary His candidate; near β-phosphate, weak broad retention. |
| UspG S128 | 396/415 | 31.0 | 33.1 | 11.6 | OG | 9.40 | 94.9 | Lower-conservation hydroxyl comparator. |
| UspG D10 | 390/415 | 97.7 | — | 93.1 | OD2 | 6.16 | 105.1 | Conserved carboxylate; distinct possible linkage or binding role. |
| YdaA T284 | 781/787 | 97.4 | 97.4 | 90.8 | OG1 | 4.55 | 122.6 | First hydroxyl candidate; contacts a γ-phosphate oxygen. |
| YdaA T270 | 781/787 | 84.9 | 90.7 | 71.0 | OG1 | 8.48 | 71.0 | Conserved hydroxyl; buried in the model. |
| YdaA S277 | 782/787 | 90.5 | 94.7 | 27.3 | OG | 11.33 | 30.7 | Conserved hydroxyl in an unresolved crystal loop; low model confidence. |
| YdaA H169 | 787/787 | 87.3 | — | 29.6 | NE2 | 5.40 | 58.6 | Nearest conserved His; non-in-line phosphate contact. |
| YdaA H225 | 787/787 | 92.9 | — | 49.3 | NE2 | 12.08 | 114.6 | Conserved His, distant and buried; lower priority. |
| YdaA H244 | 787/787 | 76.6 | — | 76.0 | ND1 | 19.91 | 147.7 | Broadly retained His, distant; lower priority. |
| YdaA C294 | 775/787 | 95.8 | — | 72.2 | SG | 17.16 | 146.5 | Conserved thiol, distant and buried; lower priority. |
| YdaA E286 | 781/787 | 100.0 | — | 93.9 | OE1 | 9.47 | 87.2 | Conserved carboxylate; distinct possible linkage or binding role. |
| YnaF S127 | 560/566 | 82.2 | 84.0 | 83.8 | OG | 5.25 | 99.8 | Corresponds to UspG S125; natural ATP-binding comparison. |
| YnaF H135 | 552/566 | 76.3 | — | 45.4 | NE2 | 14.46 | 124.4 | Known chloride-interface role; conservation has another explanation. |

Broad retention denotes Ser+Thr for Ser/Thr rows and exact residue retention for all other rows. The em dash marks a non-hydroxyl candidate. Known/N gives the unweighted count of known residues at the site and the number of sequences in the retained pool. Conservation can reflect nucleotide recognition, packing or metal coordination. Acceptor assignment requires direct identification of a covalent adduct.

**Figure S1**

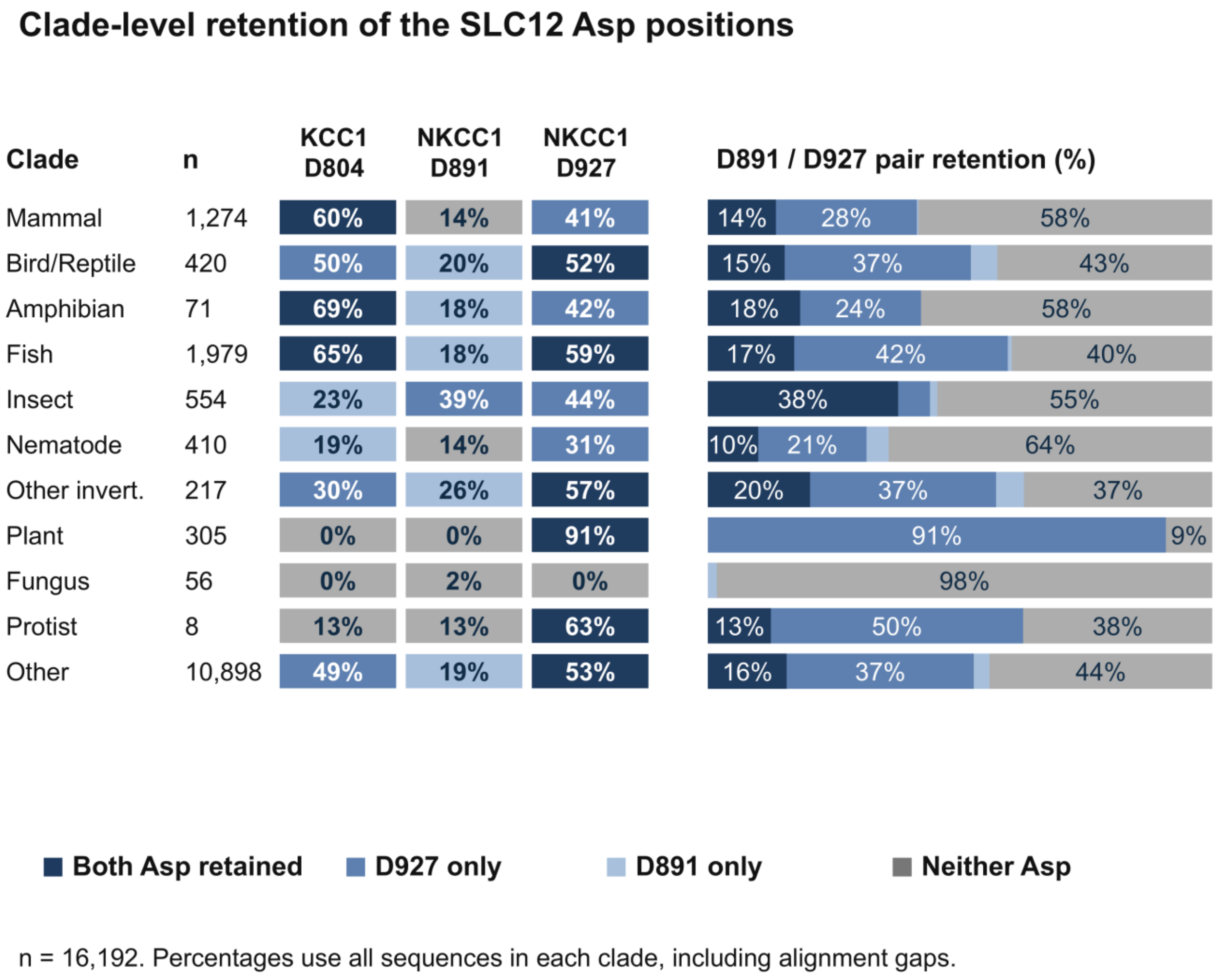


**Figure S1.** Clade-level retention of three SLC12 Asp positions. Cells show strict Asp frequencies at the alignment columns corresponding to KCC1 D804, NKCC1 D891 and NKCC1 D927. Stacked bars show both Asp residues retained, D927 only, D891 only or neither retained. Percentages use all sequences within each keyword-assigned clade, including alignment gaps (total n = 16,192). Insects retain both Asp residues in 37.7% of sequences; plants retain D927 alone in 90.8%; fungi retain neither in 98.2%. “Other” contains 10,898 sequences. The groups reflect the keyword assignments and sampled paralog composition.

**Figure S2**

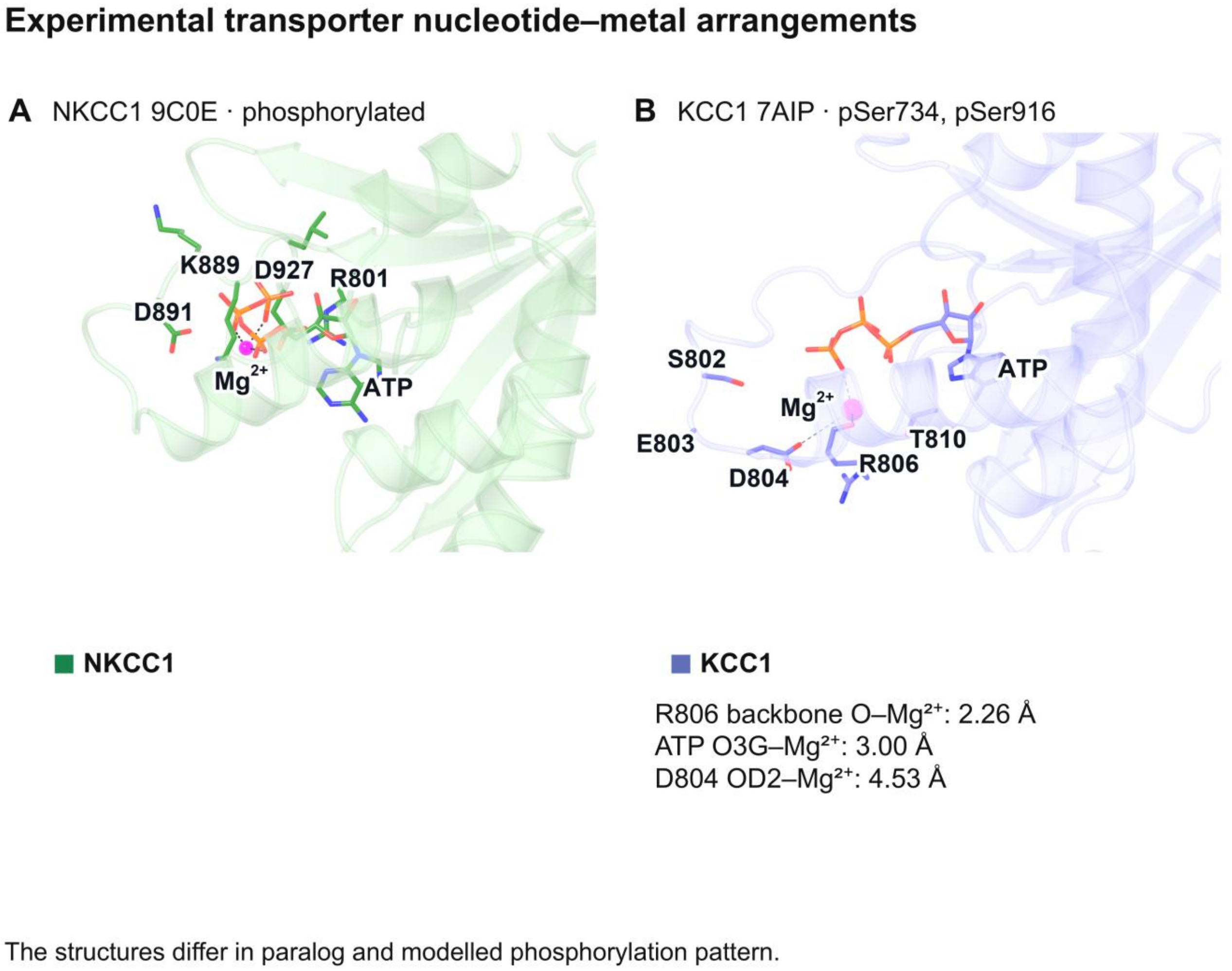


**Figure S2.** Experimental metal sites in phosphorylated NKCC1 9C0E (A) and KCC1 7AIP (B) (11, 12). The structures use the same eight-atom ribose/Pα frame and camera. KCC1 Arg806 backbone O lies 2.26 Å from its modelled $Mg^{2+}$ and is its only ligand within 2.6 Å; the nearest ATP oxygen is 3.0 Å away. In 9C0E chain A, the $Mg^{2+}$ contacts ATP oxygens at 1.80–2.51 Å and has no protein ligand. 7AIP carries modelled phosphoserines at S734 and S916, and 9C0E phosphothreonines at T212, T217 and T230, so the comparison changes both paralog and phosphorylation pattern and does not isolate a phosphorylation-induced motion.

**Figure S3**

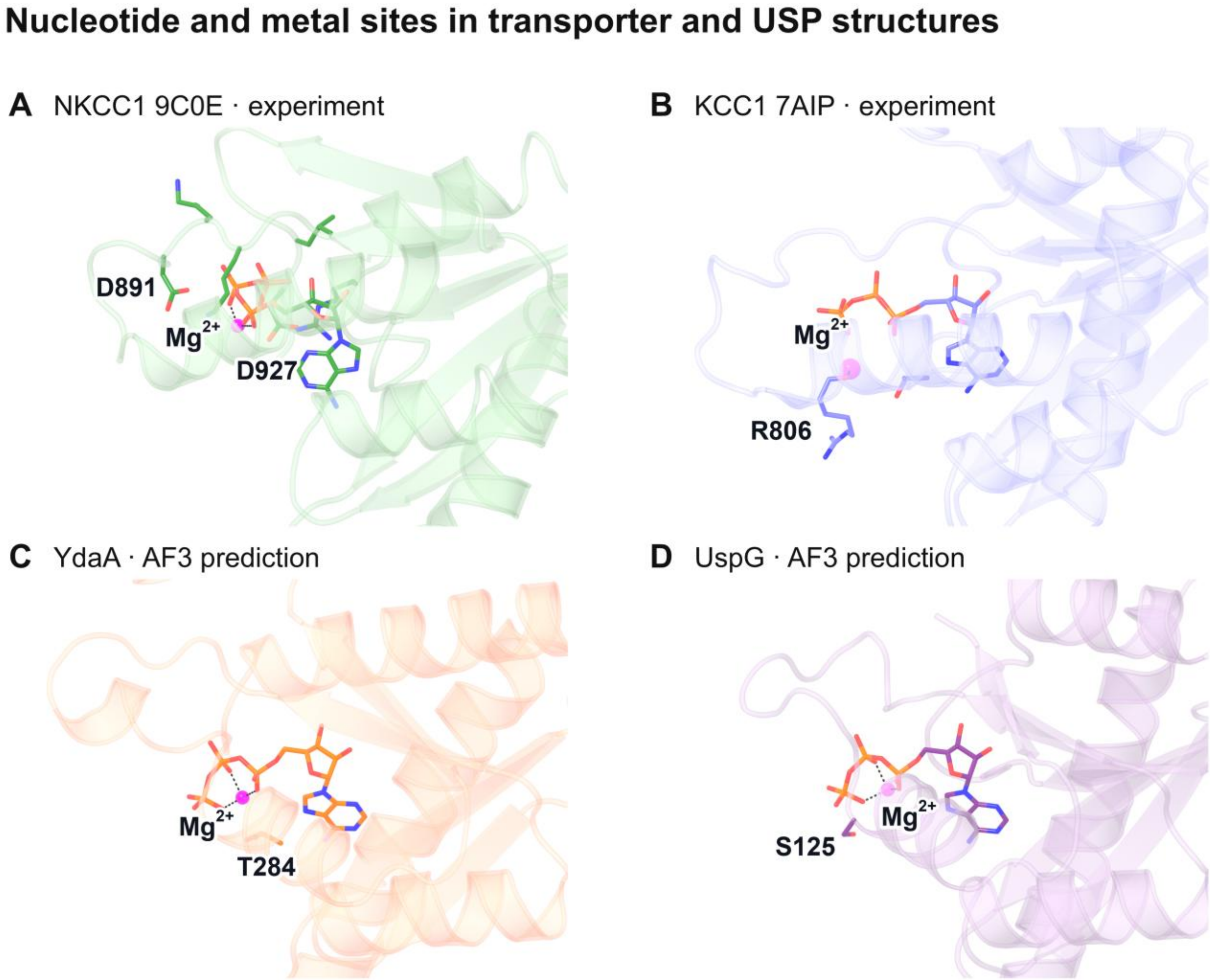


**Figure S3.** Nucleotide and metal sites in transporter and USP structures. NKCC1 9C0E (A) and KCC1 7AIP (B) are experimental structures (11, 12). YdaA (C) and natural E. coli UspG (D) show AlphaFold 3 ATP/$Mg^{2+}$ predictions: YdaA one-Mg model 0 and UspG dimer/two-Mg model 0, chain A pocket. All panels share an eight-atom ribose/Pα frame and camera. The KCC1 Arg806 backbone-oxygen contact is modelled in the experimental structure; the USP ligands are predicted. The representative USP panels contain one $Mg^{2+}$ per ATP. The additional-metal predictions are discussed in the SI text.

### Figure S4

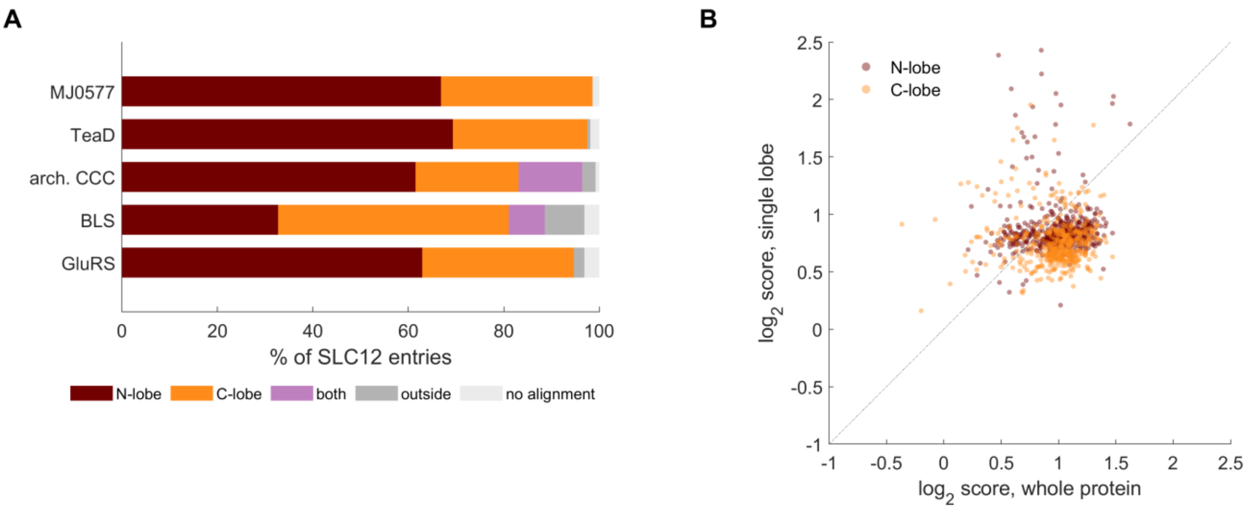


**Figure S4.** Lobe mixing in the whole-protein DALI survey. (A) Lobe of the SLC12 CTD that each reference aligned to, as a percentage of the 639 SLC12 entries. The archaeal CCC and β-lactam synthetase references align to both lobes in some entries. (B) Score of each SLC12 lobe from the domain-level survey against the whole-protein score of the same protein, on log2 axes. The dotted line is equality.

### Figure S5

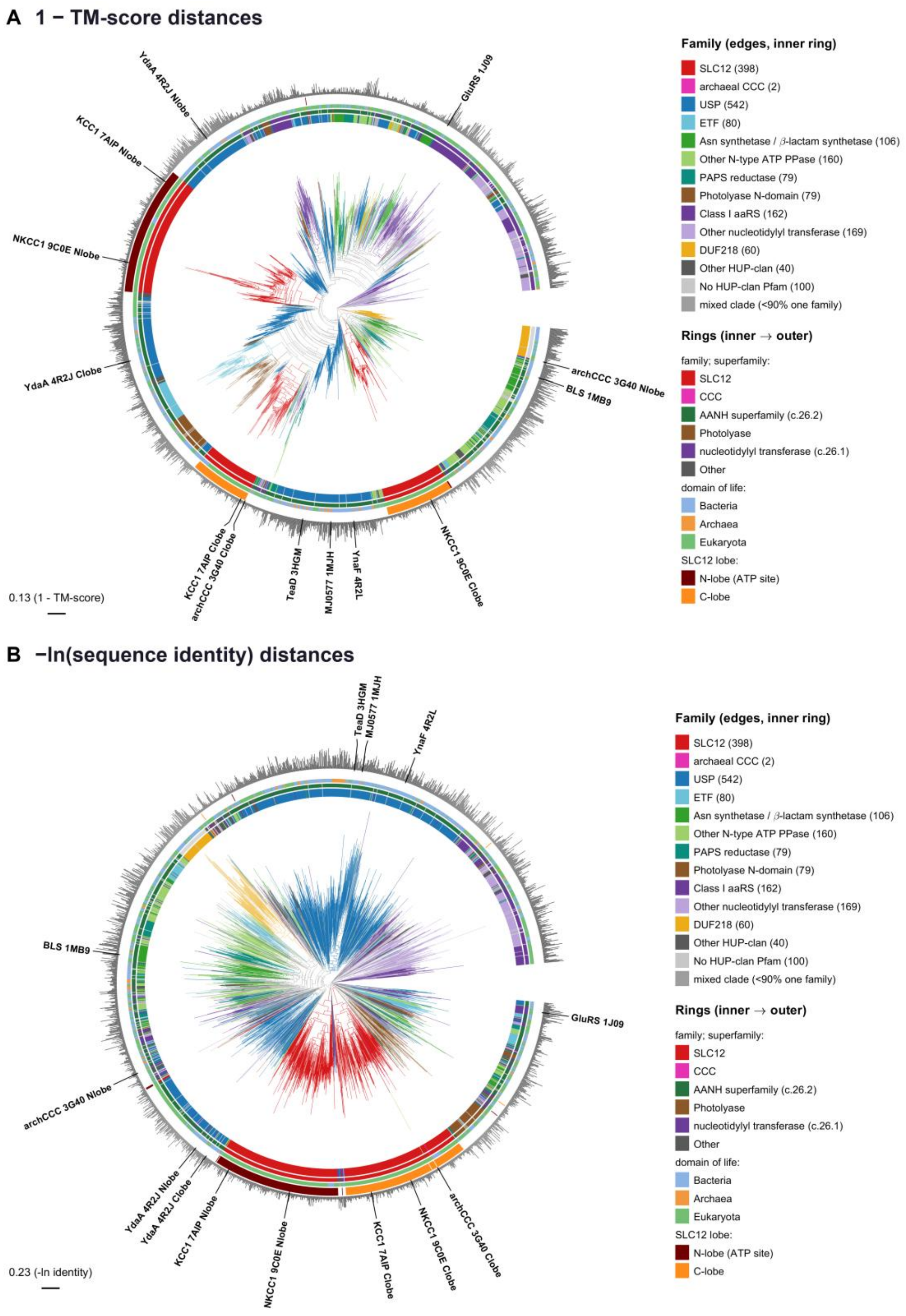


**Figure S5.** Structural trees from two further distance measures. (A) 1 − mean TM-score. (B) −ln(sequence identity) over the Foldseek alignment. Colours and rings follow Figure 4. Both trees recover the SLC12 N-lobe clade and place the archaeal CCC C-lobe with the SLC12 C-lobes; the TM-score tree splits the C-lobes into two clades (Table S4).

#### Table S3 Median DALI Z to each reference, by domain group

| Domain group | n | MJ0577 | TeaD | CCC N-lobe | CCC C-lobe | BLS | GluRS | Score |
|---|---|---|---|---|---|---|---|---|
| SLC12 N-lobe | 587 | 9.8 | 8.9 | 11.3 | 9.5 | 5.5 | 5.8 | 1.76 |
| SLC12 C-lobe | 520 | 7.8 | 6.9 | 5.6 | 9.9 | 4.9 | 5.2 | 1.66 |
| USP | 13,066 | 16.4 | 16.6 | 9.1 | 10.2 | 6.2 | 5.8 | 2.46 |
| EC-annotated | 18,665 | 7.3 | 6.9 | 6.4 | 6.3 | 5.0 | 4.8 | 1.22 |
| Other/no EC | 13,529 | 8.0 | 7.8 | 7.0 | 6.6 | 5.3 | 4.9 | 1.45 |

Z values are medians over the domains of each group in the domain-level survey. The score is the mean of the MJ0577, TeaD and higher-scoring CCC lobe values divided by the mean of the BLS and GluRS values, taken per domain. Domains without an alignment to a reference contribute Z = 0. The score median is over the domains with a defined score (577, 493, 12,947, 18,326 and 13,035).

#### Table S4 SLC12 lobe clades across three structural distance measures

| Distance | SLC12 N-lobe clade | SLC12 C-lobe clade | Archaeal CCC C-lobe | Bacterial SLC12 lobes |
|---|---|---|---|---|
| 1 − LDDT | 199 tips, 199 of 202 N-lobes | 196 tips, 195 of 196 C-lobes | sister to the C-lobe clade (+1 tip) | 14/14 N, 13/13 C |
| 1 − TM-score | 195 tips, 195 of 202 N-lobes | two clades (104, 92 tips) | sister to the 92-tip clade | 14/14 N, 13/13 C |
| −ln identity | 197 tips, 196 of 202 N-lobes | 193 tips, 191 of 196 C-lobes | in the C-lobe clade | 14/14 N, 13/13 C |

Each clade is the smallest one holding at least 95% of the lobes of that type. Tip counts include the experimental lobes. The 1 − LDDT tree is shown in Figure 4 and the other two in Figure S5. For the TM-score C-lobes no clade meets this rule; the two largest clades made only of C-lobes are given.

### References


1. Weber A, Jung K (2006). Biochemical Properties of UspG, a Universal Stress Protein of Escherichia coli. Biochemistry 45:1620–1628. https://doi.org/10.1021/bi051301u

2. Bangera M, Panigrahi R, Sagurthi SR, Savithri HS, Murthy MRN (2015). Structural and functional analysis of two universal stress proteins YdaA and YnaF from Salmonella typhimurium: possible roles in microbial stress tolerance. J Struct Biol 189:238–250. https://doi.org/10.1016/j.jsb.2015.01.003

3. Xu Y, Guo J, Jin X, Kim JS, Ji Y, Fan S, et al. (2016). Crystal structure and functional implications of the tandem-type universal stress protein UspE from Escherichia coli. BMC Struct Biol 16:3. https://doi.org/10.1186/s12900-016-0053-9

4. Henikoff S, Henikoff JG (1994). Position-based sequence weights. J Mol Biol 243:574–578. https://doi.org/10.1016/0022-2836(94)90032-9

5. Abramson J, Adler J, Dunger J, Evans R, Green T, Pritzel A, et al. (2024). Accurate structure prediction of biomolecular interactions with AlphaFold 3. Nature 630:493–500. https://doi.org/10.1038/s41586-024-07487-w

6. Gonzalez A, Pon A, Servage K, Pawłowski K, Han Y, Sreelatha A (2025). A repurposed AMP binding domain reveals mitochondrial protein AMPylation as a regulator of cellular metabolism. Nat Commun 16:7863. https://doi.org/10.1038/s41467-025-63014-z

7. Zhang Z, Patel R, Luo ZQ, Das C (2025). Cryo-EM Detection of AMPylated Histidine Implies Covalent Catalysis in AMPylation Mediated by a Bacterial Effector. J Mol Biol 437:168917. https://doi.org/10.1016/j.jmb.2024.168917

8. He C, Li C, Liu Y, Chen TT, Li C, Chu X, et al. (2025). Modulation of host ATP levels by secreted bacterial effectors. Nat Commun 16:4675. https://doi.org/10.1038/s41467-025-60046-3

9. Passaro S, Corso G, Wohlwend J, Reveiz M, Thaler S, Somnath VR, et al. (2025). Boltz-2: Towards Accurate and Efficient Binding Affinity Prediction. bioRxiv. https://doi.org/10.1101/2025.06.14.659707

10. Hori T, Moriwaki Y, Ishitani R (2026). Distance-Restraint-Guided Diffusion Models for Sampling Protein Conformational Changes and Ligand Dissociation Pathways. J Chem Theory Comput 22:5259–5270. https://doi.org/10.1021/acs.jctc.6c00199

11. Zhao Y, Vidossich P, Forbush B, Ma J, Rinehart J, De Vivo M, et al. (2025). Structural basis for human NKCC1 inhibition by loop diuretic drugs. EMBO J 44:1540–1562. https://doi.org/10.1038/s44318-025-00368-6

12. Chi G, Ebenhoch R, Man H, Tang H, Tremblay LE, Reggiano G, et al. (2021). Phospho-regulation, nucleotide binding and ion access control in potassium-chloride cotransporters. EMBO J 40:e107294. https://doi.org/10.15252/embj.2020107294
